# Meiosis-specific functions of kinesin motors in cohesin removal and maintenance of chromosome integrity in budding yeast

**DOI:** 10.1101/692145

**Authors:** Priyanka Mittal, Hemant Kumar Prajapati, Komal Ghule, Santanu K. Ghosh

## Abstract

Kinesin motors provide the molecular forces at the kinetochore-microtubule interface and along the spindle to control chromosome segregation. During meiosis with the two rounds of microtubule assembly-disassembly, the roles of motor proteins remain unexplored. We observed that in contrast to mitosis Cin8 (kinesin 5) and Kip3 (kinesin 8) together are indispensable in meiosis. Examining the meiosis in *cin8*∆ *kip3*∆ cells, we detected chromosome breakage in the meiosis II cells. The double mutant exhibits delay in the cohesin removal and spindle elongation during anaphase I. Consequently, some cells abrogate meiosis II and form dyads while some, as they progress through meiosis II, cause defect in chromosome integrity. We believe that in the latter cells, an imbalance of spindle mediated force and simultaneous persistent cohesin on the chromosomes cause their breakage. We provide evidence that tension generated by Cin8 and Kip3 through microtubule cross-linking is essential for signaling efficient cohesin removal and maintenance of chromosome integrity during meiosis.

**Summary:** Molecular motors generate forces that facilitate chromosome segregation. Unlike mitosis, in meiosis, two times chromosome segregation occur with twice microtubule assembly/disassembly. This work reports that the motor mediated forces are crucial for cohesin removal in meiosis and thus maintain genome integrity.

## Introduction

Meiotic chromosome segregation comprises of certain unique events unlike in mitosis. In budding yeast these events include late assembly of the mature kinetochore competent to connect the microtubules; pairing of the homologs; mono-orientation of the sister kinetochores with respect to the spindle pole in metaphase I; a step wise dissolution of cohesin from the chromatin; two rounds of chromosome segregation with spindle assembly and dis-assembly and a partial dephosphorylation of the CDK substrates adequate for spindle disassembly at meiosis I but not for DNA replication. Further, while dephosphorylation of the CDK substrates by Cdc14 phosphatase is essential for cell cycle exit both in mitosis and meiosis, FEAR-released Cdc14 appears to be dispensable for the same in mitosis but not in meiosis as it is required to exit from meiosis I (Buonomo et al., 2003; Marston et al., 2003; Yellman and Roeder, 2015). Nevertheless, FEAR-released Cdc14 has significant roles both in mitosis and meiosis for coherent segregation of all the chromosomal loci, and for the stability and proper orientation of the microtubule spindle (D’Amours and Amon, 2004; Khmelinskii and Schiebel, 2008; Lee and Amon, 2003; Marston et al., 2003; Roccuzzo et al., 2015).

Irrespective of the type of cell cycle, the formation of a microtubule-based spindle and movement of the chromosomes along the spindle being attached to the microtubule play a pivotal role during chromosome segregation. The occurrence of these events relies on the polymerization-depolymerization property of the microtubules which is facilitated by the functions of several microtubule-associated proteins (MAPs) and microtubule based motors (Barton and Goldstein, 1996). In *S. cerevisiae* four nuclear motors of kinesin superfamily (Kinesin-related Proteins, KRPs) namely Cin8, Kip1, Kip3 and Kar3, have essential roles in chromosome segregation (Gupta Jr et al., 2006; Roof et al., 1992; Saunders and Hoyt, 1992; Saunders et al., 1995; Tytell and Sorger, 2006). However, due to functional redundancy among these motors, they are non-essential for growth (Cottingham et al., 1999; DeZwaan et al., 1997).

Cin8 and Kip1 belong to BimC or kinesin-5 family of proteins, where the motor domain is at the amino terminal end of the protein and the motor movement is directed towards the plus end of the microtubule (Dagenbach and Endow, 2004; Hoyt et al., 1992a). Cin8 and Kip1 form homo-tetramers and their plus-end directed functions are to extend the spindle by pushing the poles apart and to maintain the kinetochores in the clustered form through cross-linking of antiparallel and parallel microtubules, respectively (Gordon and Roof, 1999; Hildebrandt et al., 2006; Kashlna et al., 1996; Mayr et al., 2011; Rizk et al., 2014). Later, through in vitro assays, minus-end directed movement of both of these single motors have been identified when they work singly on individual microtubule (Fridman et al., 2013; Gerson-Gurwitz et al., 2011; Roostalu et al., 2011; Shapira and Gheber, 2016). Recently, it has shown under in vivo condition that Cin8 clusters at the minus end and SPB during the early stage of mitosis for capturing the microtubules emanating from opposite SPBs that facilitates bi-polar spindle formation (Shapira et al., 2017). However, the implication of Kip1 minus end directed movement has not been explored. In addition to the cross-linking function, Cin8 and to a lesser extent Kip1 can also depolymerize kMT (kinetochore microtubule) in a length dependent manner which is believed to be essential for congression of the chromosomes (Gardner et al., 2008). The regulation of Cin8 and Kip1 functions depends on the phosphorylation status of these proteins where their phosphorylation by Cdk1 during early mitosis mediates the SPB separation (Chee and Haase, 2010). In metaphase, Cin8 and Kip1 are localized at the centromeres and along the microtubule length (Tytell and Sorger, 2006). Since phosphorylation of Cin8 inhibits its association to the microtubules (Goldstein et al., 2017), following metaphase to anaphase transition dephosphorylation of Cin8 by PP2A^Cdc55^ and Cdc14 results in its accumulation near the spindle poles and at the spindle midzone which is crucial for spindle elongation (Avunie-Masala et al., 2011; Goldstein et al., 2018). However, it is not known if similar dephosphorylation also occurs in Kip1. During early anaphase, APC^Cdc20^ degrades Kip1 (Gordon and Roof, 2001) whereas Cin8 is degraded during late anaphase by APC^Cdh1^ (Hildebrandt and Hoyt, 2001). On the other hand the primary function of Kip3 motor, belonging to either kinesin-8 or −13 family of proteins, is the depolymerization of microtubules plus ends by a mechanism similar to kinesin-13 motors (Friel and Howard, 2011; Gupta Jr et al., 2006; Gupta et al., 2006; Hunter et al., 2003). This function of Kip3 driven by its neck domain with depolymerase activity (Su et al., 2013a) has a role in the movement of chromosomes during anaphase (Tang and Toda, 2015; Tytell and Sorger, 2006). However, Kip3 also slides and clusters the microtubules by cross-linking antiparallel and parallel microtubules, respectively through its tail domain (Su et al., 2011a). Although, the cross-linking function of Kip3 is trivial as compared to kinesin-5 proteins owing to it’s intrinsic structural ability to form homo-dimer but not homotetramer observed in kinesin-5 motors (Gordon and Roof, 1999; Hildebrandt et al., 2006; Kashlna et al., 1996; Mayr et al., 2011; Rizk et al., 2014). Kip3 activity appears to be regulated spatially and temporally based on the length of the spindle and the exact localization of the motor. On a short spindle, it helps in clustering and alignment of the kinetochores by cross-linking of the parallel microtubules and depolymerase activity at the plus ends. During increase in the spindle length Kip3 cross-links and slides the anti-parallel interpolar microtubules. Finally when the spindle reaches to its maximum length, Kip3 localize at the plus-ends and by its depolymerization activity causes spindle disassembly (Rizk et al., 2014; Su et al., 2013a). Kar3 (a minus end-directed kinesin-14 family of protein) is a second microtubule depolymerizer present in the cell and is functionally antagonistic to Cin8/Kip1 spindle elongation activity. Kar3 pulls two spindle poles together, therefore the spindle collapse observed in the absence of both Cin8 and Kip1 can be suppressed by reducing the activity of Kar3 (Hoyt et al., 1993). Additionally, Kar3 appears to promote kinetochore-microtubule attachment as in mitosis it is found to occupy a subset of kinetochores on which microtubule attachments are slow to form (Tytell and Sorger, 2006).

Several groups as described above have elucidated the functions of the nuclear kinesin motors in chromosome segregation in mitosis. Given the mechanistic uniqueness in chromosome segregation in meiosis as outlined above, it is intriguing to investigate their functions during this cell cycle. However, *KAR3* mutant was found to be arrested at prophase I (Bascom-Slack and Dawson, 1997; Shanks et al., 2001) and hence makes it difficult to analyze the meiotic events in lack of Kar3. Therefore, in this study, we focused on elucidating the functions of the three motors, Cin8, Kip1 and Kip3 in meiosis. Using knockout mutants, we observed that these motors are required for homolog pairing. Strikingly, we noticed cells with loss of both Cin8 and Kip3 harbor chromosome breakage. Further investigation argues for a defect in Rec8-cohesin removal from the chromatin in these cells. We propose that the conditions in the absence of Cin8 and Kip3 perhaps create an imbalance between the microtubule mediated force and the resisting force by the persistent cohesin, which may lead to the chromosome breakage. From our findings, we suggest that the tension generated by the cross-linking activity of Cin8 and Kip3 is crucial to signal the cells for the cohesin cleavage. Thus, our study reveals significant roles of kinesin motors in meiosis and hints towards essentiality of these proteins in suppressing aneuploidy during gametogenesis.

## Results

### The motors are required for faithful meiosis

As the first set of experiments, we compared the spore viability, a readout for faithful meiosis, between the wild-type and the individual motor mutants. Given that there are functional redundancies among the motors, we observed a marginal decrease in spore viability in *kip1*Δ and *kip3*Δ (approximately 89 and 92%, respectively). However, *cin8*∆ showed around 65% reduction in spore viability suggesting this protein is more significant in meiosis (Fig. 1A). It is expected that the pace of meiotic progression can be slowed down if there is any perturbation in meiosis. To test this wild-type and the mutant strains were released into synchronized meiosis. Consistent with the spore viability data we observed that *cin8*∆ showed a delay at metaphase I compared to the wild-type, *kip1*∆ and *kip3*∆ mutants (Fig. 1B, ii) suggesting some defect is occurring during early meiotic events in the absence of Cin8 and perhaps due to functional redundancy the defect is not apparent in *kip1*∆ and *kip3*∆ mutants. To investigate if the defect causes chromosomes to mis-segregate we marked both the CenV homologs with TetO/TetR-GFP system (see materials and methods) and observed their distribution at the end of meiosis. Following faithful meiosis, a tetra-nucleate would show one GFP dot at each nucleus (Fig. 1D; Type I). However, four GFP dots in three, two nuclei (Type II and III) or in one nucleus accounts for chromosome mis-segregation. The meiotic induction, unless otherwise mentioned, was carried out at 33°C, as the phenotype of loss of Cin8 becomes aggravated at a higher temperature (Hildebrandt and Hoyt, 2000; Saunders et al., 1997; Tytell and Sorger, 2006). We observed around 50, 22 and 17% (Type II and Type III) of chromosome mis-segregation in *cin8*∆, *kip1*∆ and *kip3*∆ cells, respectively (Fig. 1D) suggesting spore viability defect is probably due to the generation of aneuploids. As in *cin8*∆, the delay in the cell cycle occurs in metaphase I, we presumed that at least some defects might be occurring during the preceding events of chromosome segregation that include chromatid cohesion, homolog pairing and sister chromatid mono-orientation. To investigate the cohesion between the sisters and the orientation of their spindle attachment, both the sisters of one homolog were marked with TetO/TetR-GFP system. In the metaphase I arrested cells, a defect in sister chromatid mono-orientation would appear as two GFP dots. On the other hand, non-cohesed sisters in the cycling cells would produce bi-nucleates with one GFP dot in each of the nucleus. However, we failed to detect any defect either in sister chromatid monoorientation or in their cohesion (Fig. S1A-B). Although not for Cin8, Kip1 or Kip3, a role of Kar3 in sister chromatid cohesion in mitosis has been reported before (Mayer et al., 2004).

**Figure 1:**
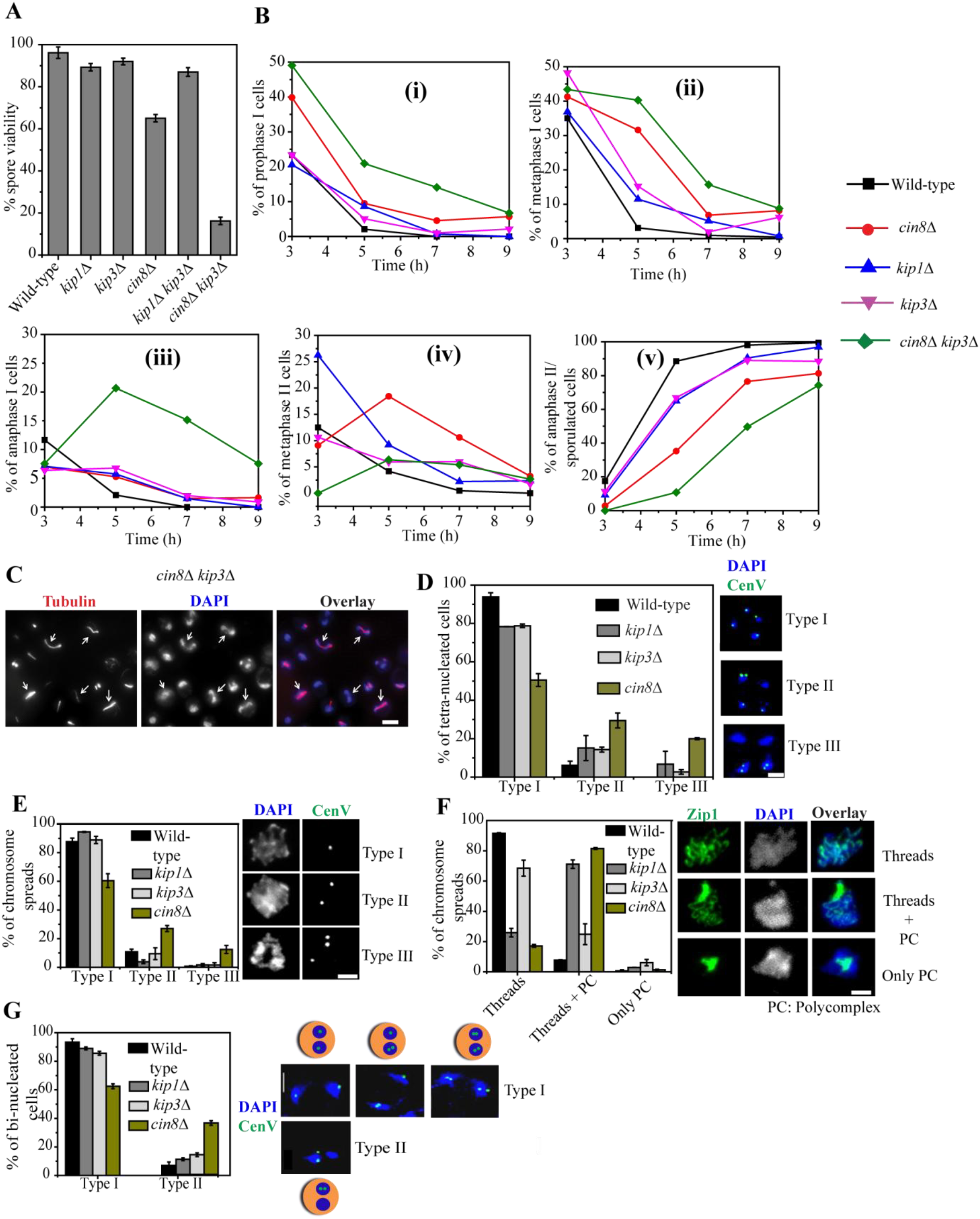
The analysis of meiosis in the motor mutants and their roles in pairing and disjunction of the homologs. (A) The spore viability of the wild-type (SGY5001; n = 40), *cin8*∆ (SGY315; n = 60), *kip1*∆ (SGY317; n = 107), *kip3*∆ (SGY314; n = 91), *kip1*∆ *kip3*∆ (SGY5104; n = 54), and *cin8*∆ *kip3*∆ (SGY5089; n = 117) cells were analyzed after inducing meiosis at 30°C. ‘n’ represents the number of tetrads dissected. (B) The indicated strains were induced for synchronized meiosis and analyzed for meiotic progression. At the above mentioned time points, the fraction of cells at different stages of meiosis was determined by anti-α-tubulin staining. At least 100 cells were counted for each time point. (C) Representative image of *cin8*∆ *kip3*∆ cells showing the maximum population at anaphase I stage at 5 h of meiotic induction as determined by anti-α-tubulin and DAPI staining. Arrows represent the anaphase I cells. Bar, 5 µm. (D) The indicated strains harboring homozygous CenV-GFP (see material and methods) were analyzed for the meiotic chromosome segregation at 33°C in the tetra-nucleated cells (n = 90-150). (E-F) The wild-type (SGY263), *cin8*∆ (SGY5197), *kip1*∆ (SGY5347), and *kip3*∆ (SGY5190) cells arrested at prophase I by Ndt80 depletion and harboring homozygous CenV-GFP were analyzed by chromosome spreads for the number of GFP dots and Zip1 localization in (E) and (F), respectively (n = 150-220). 1 or 2 or > 2 GFP dot(s) were scored as paired or unpaired or unpaired with non-cohesed sister chromatids, respectively. (G) Strains as mentioned in (A) were analyzed at the bi-nucleated stage for the disjunction of CenV homologs (n = 50 – 110). Type I and II indicate disjunction and non-disjunction of the homologs, respectively. Error bars represent the standard deviation from the mean values obtained from at three independent experiments. Bar, 2 µm.

However, we observed an increased defect in homolog pairing in *cin8*∆ when both the CenV homologs were marked with GFP (Fig. 1E; Type II and Type III). Consistent with this we observed a higher percentage of mis-localization (polycomplex formation) of Zip1, a component of the synaptonemal complex (SC) that reinforces pairing (Fig. 1F). A similar result was also obtained before where homologs fail to synapse in the absence of Kar3 (Bascom-Slack and Dawson, 1997). Following dis-assembly of SC, Zip1 is maintained at the centromeres until the proper bipolar attachment of the homologs is achieved (Gladstone et al., 2009). As the Zip1 localization was compromised in *cin8*∆ or *kip1*∆ mutant, we examined the homolog biorientation in the motor mutants where both the CenV homologs were marked with GFP. About 37% of the bi-nucleated cells in *cin8*∆ showed homolog non-disjunction compared to only 7% in the wild-type (Fig. 1G; Type II).While in *kip3*∆ and *kip1*∆, the population exhibiting such defect was relatively less (14% and 11%, respectively, Fig.1G). Similar results were observed in *zip1∆ mad2*∆ double mutant where about 45% of cells showed homolog non-disjunction (Gladstone et al., 2009) which is comparable to the population obtained in *cin8*∆ in our study. Above results suggest that the absence of the motor proteins, specially Cin8, can affect homolog pairing in meiosis.

### Meiosis is profoundly compromised in *cin8∆ kip3*∆ double mutant

Since Cin8, Kip1 and Kip3 share overlapping functions in microtubule cross-linking and depolymerization (Gardner et al., 2008; Gupta et al., 2006; Hoyt et al., 1992a; Saunders and Hoyt, 1992; Su et al., 2013a; Tytell and Sorger, 2006; Winey and Bloom, 2012), we argued that their functions cannot be properly revealed studying only the single mutants. Therefore, we generated two possible viable double mutants *kip1*∆ *kip3*∆, and *cin8*∆ *kip3*∆ as *cin8*∆ *kip1*∆ has been reported as inviable (Gordon and Roof, 1999; Hoyt et al., 1992a; Kapitein et al., 2005; Roof et al., 1992). Albeit, the sporulation efficiencies of *cin8*∆, *kip3*∆ and *cin8*∆ *kip3*∆ strains were similar (63%, 85% and 70%, respectively; Fig 2A) after 12 h of sporulation induction, strikingly we observed a precipitous drop (approximately 16%) in spore viability of the double mutant compared to *cin8*∆ and *kip3*∆ single mutants (Fig. 1A). To further investigate the probable roles of Cin8 and Kip3 together in meiosis, we followed meiotic progression in the wild-type and motor mutants and noticed that in comparison to the wild-type or *cin8*∆ or *kip3*∆, *cin8*∆ *kip3*∆ double mutant proceeded through meiosis more slowly and the majority of the cells were arrested transiently at anaphase I with one spindle and improper disjunction of nuclei (Fig.1B, iii-C).

**Figure 2:**
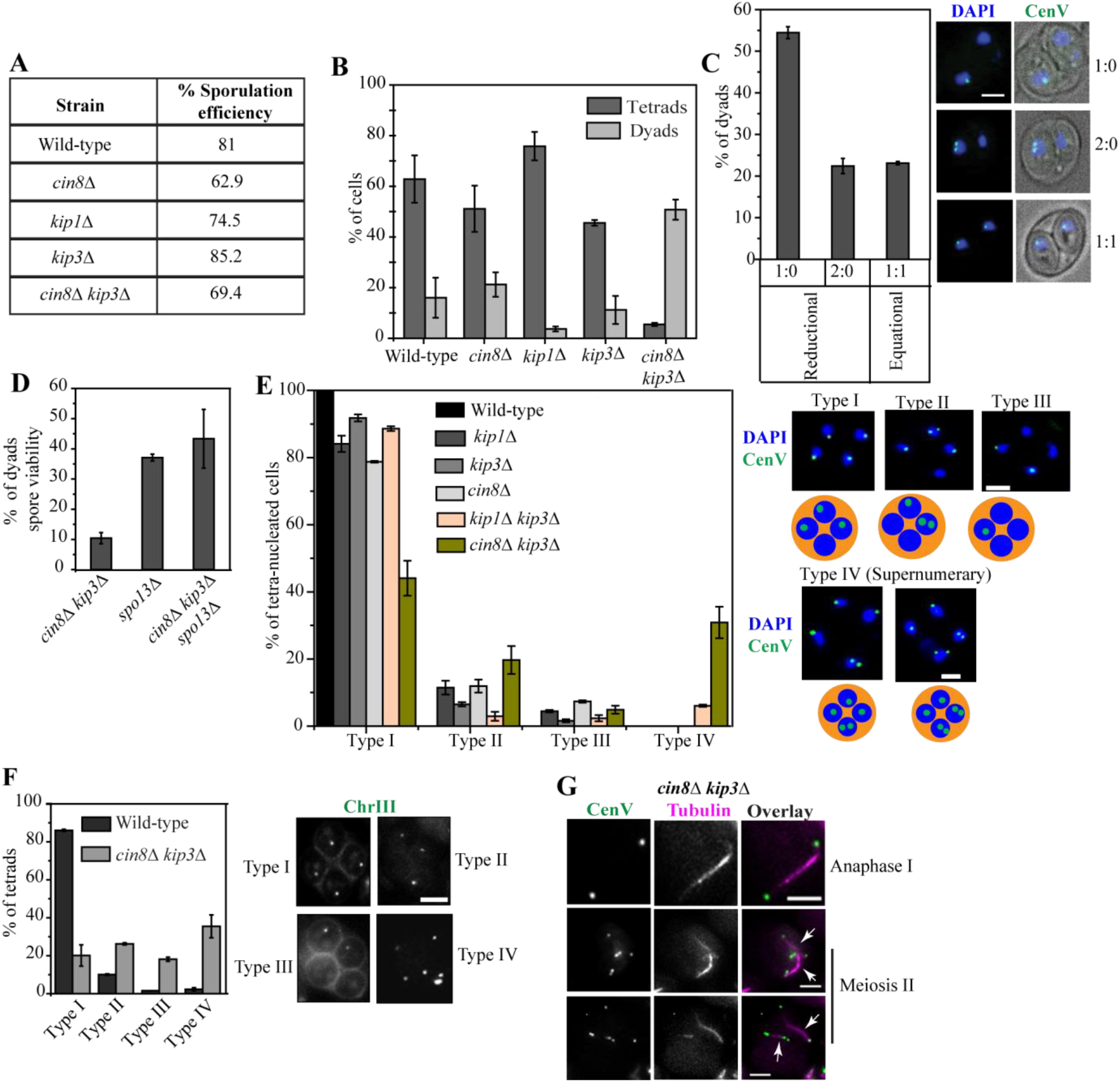
Meiosis in *cin8*∆ *kip3*∆ cells results in frequent formation of dyads with aneuploid spores and also generates aneuploid tetra-nucleated cells showing supernumerary GFP foci of the marked chromosomes. (A-B) The wild-type (SGY5001), *cin8*∆ (SGY315), *kip1*∆ (SGY317), *kip3*∆ (SGY314), and *cin8*∆ *kip3*∆ (SGY5089) cells were analyzed for the (A) sporulation efficiency and for the (B) formation of dyads and tetrads following 12 h of meiotic induction (n = 195-351). The maximum population of the *cin8*∆ *kip3*∆ sporulated cells form dyads with a small population of tetrads. ‘n’ represents the total number of sporulated cells scored for the analysis. (C) The percentage of dyads with the one GFP dot each in two spores (1:1) and one or two GFP dot(s) in one spore (1:0 or 2:0, respectively) were determined in *cin8*∆ *kip3*∆ (SGY5154; n = 136) harboring heterozygous CenV-GFP. Bar, 2 µm. (D) The spore viability of the dyads formed in *cin8∆ kip3*∆ (SGY5089), *spo13*∆ (SGY402), and *cin8∆ kip3*∆ *spo13*∆ (SGY5442) following meiosis at 30°C. For each strain, 60 dyads were dissected for the viability estimation. (E) The tetra-nucleated cells from the indicated strains harboring homozygous CenV-GFP were analyzed for the meiotic chromosome segregation at 30°C (n = 100-309). ‘n’ represents the number of tetra-nucleated cells scored for chromosome segregation. (F) The tetrads from the the wild-type (SGY5407; n = 129) and *cin8*∆ *kip3*∆ (SGY5329; n = 119) harboring homozygous ChrIII-GFP marked at *LEU2* locus 22 kb away from the centromere were analyzed as in (E). (G) The supernumerary CenV-GFP foci were observed in only those *cin8*∆ *kip3*∆ cells (SGY5385) that are in meiosis II as judged by the presence of two spindles (marked by arrows). Error bars represent the standard deviation from the mean values obtained from three independent experiments.

Given that Cin8 and Kip3 can cross-link and slide the antiparallel microtubules causing spindle elongation, our results indicate that the cells lacking both Cin8 and Kip3 cause slow spindle elongation and defects in chromosome disjunction during meiosis I that might be responsible for a delay in spindle disassembly and completion of meiosis I. Furthermore, due to this delay around 50% of the *cin8*∆ *kip3*∆ cells biochemically proceeded to meiosis II without completing meiosis I and produced dyads (Fig. 2B). Inability to complete meiosis I due to defect in spindle elongation but proceeding to meiosis II and generation of the dyads with two diploid spores are the hallmarks of the FEAR mutants (Buonomo et al., 2003; Kamieniecki et al., 2000; Marston et al., 2003; Zeng and Saunders, 2000). Additionally, similar to the FEAR mutants (Marston et al., 2003), in *cin8*∆ *kip3*∆ cells, we mostly observed reductional segregation of chromosomes in the dyads as both the heterozygously tagged CenV-GFP dots (sister chromatids) were found in one spore in 76% of the dyads (Fig. 2C). However, co-segregation of the sister chromatids per se does not imply the abrogation of meiosis II in *cin8*∆ *kip3*∆ cells since in many dyads we observed stained nuclei that were not included into the spores suggesting massive mis-segregation has occurred during meiosis II as well. Consequently, the viability of the dyad spores obtained from *cin8*∆ *kip3*∆ cells was extremely poor (10%; Fig. 2D).

We believe that the phenotypes of *cin8*∆ *kip3*∆ cells are similar to the FEAR mutants as the FEAR-released Cdc14 phosphatase promotes spindle elongation through dephosphorylation of Cin8 that facilitates its binding to the spindles and sliding of the anti-parallel microtubules (Avunie-Masala et al., 2011; Roccuzzo et al., 2015). However, removal of Cin8 alone did not exhibit as severe phenotype as the FEAR mutants due to functional redundancy in spindle elongation between Cin8 and Kip1/Kip3 and due to additional functions of the FEAR network (Rock and Amon, 2009). It is also expected that the FEAR mutant-like phenotypes observed in meiotic *cin8*∆ *kip3*∆ cells will also be observed in mitosis. Since the FEAR mutants exhibit a delay in mitotic exit (Stegmeier et al., 2002), the wild-type and the motor mutants were released synchronously using α-factor into fresh YPD media to compare the pace of mitosis. As observed for the FEAR mutant *esp1-1* (Queralt et al., 2006; Stegmeier et al., 2002), we observed a delay in cell-cycle progression in *cin8*∆ and *cin8*∆ *kip3*∆ cells. Wild-type, *kip1*∆, and *kip3*∆ completed one cycle of mitosis in approximately 55 min while in *cin8*∆ and *cin8*∆ *kip3*∆ it was delayed (around 75 min) as shown by dotted and dashed line, respectively in figure S2A. In the time window between 90 to 100 min (Fig. S2B), the wild-type, *kip1*∆, and *kip3*∆ exhibit the second peak for the metaphase cells, while *cin8*∆ demonstrates only one peak whereas in *cin8*∆ *kip3*∆ the same peak is broadened and further extended till 105 min suggesting that metaphase to anaphase transition is maximally delayed in the double mutant. Additionally, we also observed a phenotype in *cin8*∆ and *cin8*∆ *kip3*∆ mutants at an equal frequency where co-ordination between spindle elongation and the chromosome segregation was compromised. In these cells we found a persistent population of cells with elongated nucleus spanning mother and daughter buds but with a bipolar spindle in one of the buds with a length specific to that of the metaphase (Fig. S2C; Type II). Such a phenotype could be due to inability to extend the spindle but with continuous to and fro movement of the short spindle resulting nuclear elongation which has been observed before with metaphase arrested short spindle (Tanaka et al., 2007)

From the above results, it is apparent that absence of both Cin8 and Kip3 causes defects in spindle elongation, metaphase to anaphase transition and spindle disassembly both in mitosis and meiosis. However, in contrast to poor spore viability following meiosis observed in the *cin8*∆ *kip3*∆ (Fig. 1A), we failed to get any difference in viability among the wild-type, single mutants and *cin8*∆ *kip3*∆ double mutant following mitosis (Fig. S2D). This is further supported by the fact that while the pace of meiosis was found to be affected to a large extent in the *cin8*∆ *kip3*∆ double mutant with respect to the wild-type or the single mutants (Fig.1B), the mitotic growth rates were not affected to that extent (Fig. S2E). These results suggest that loss of both Cin8 and Kip3 perhaps cause some meiosis specific defects as revealed below.

### Meiotic chromosome segregation is largely perturbed in *cin8∆ kip3*∆

To examine if there are any meiotic-specific defects in *cin8*∆ *kip3*∆ double mutant we sought out to investigate the meiotic chromosome segregation under this condition. We used wild-type, and *cin8*∆ *kip3*∆, *kip1*∆ *kip3*∆ double mutants and the corresponding single mutants cells where both the homologs of chromosome V were marked with CenV-GFP. Since we observed that *cin8*∆ *kip3*∆ double mutant did not sporulate at increased 33° C temperature, meiosis induction was carried out at 30°C.We analyzed the tetra-nulceated cells to ensure both meiosis I and II have occurred. Wild-type, *kip3*∆, *kip1*∆, and *kip1*∆ *kip3*∆ cells showed mostly (100%, 92%, 84%, and 88%, respectively) four nuclei with one GFP dot in each (Fig. 2E; Type I) which was reduced in *cin8*∆ and largely in *cin8*∆ *kip3*∆ cells (79% and 44%, respectively). Type II and III categories having GFP dots in three or in two nuclei, respectively and accounting for mis-segregation of the chromosomes were found correspondingly more in the mutants. Unexpectedly, a significant population of tetra-nucleates (approximately 30%) harboring >4 (termed ‘supernumerary’) CenV*-*GFP dots were found in *cin8*∆ *kip3*∆ cells (Fig. 2E; type IV category) while a minute population of this category was observed in *kip1*∆ *kip3*∆ cells (6%). The difference observed between the double mutants can be expected since Cin8 is known to have more significant cell cycle functions than Kip1 from the mitotic study (Hoyt et al., 1992b). The supernumerary GFP dots phenotype is not specific to chromosome V since the same phenotype was also observed (approximately 35%; Type IV) in the *cin8*∆ *kip3*∆ double mutant when chromosome III was marked using LacO/LacI-GFP system at the pericentromeric region (22 kb away from CenIII; Fig. 2F). To determine the stage of the cell cycle at which these supernumerary foci start appearing, Tub1 was N-terminally tagged with CFP in the *cin8*∆ *kip3*∆ strain harboring homologous CenV-GFP. Chromosome abnormality was found only in the cells with two spindles suggesting >4 foci were generated in the cells that have passed through meiosis II (Fig. 2G). This numerical abnormality is specific to meiosis and did not occur due to any aneuploidy generated as a legacy of an error during previous mitosis since we failed to obtain >2 GFP dots before entering into meiosis (Fig. S3A) or during any stage of mitosis (Fig. S3B) in the *cin8*∆ *kip3*∆ cells.

### Chromosome breakage occurs in *cin8*∆ *kip3*∆ during meiosis II

We next sought to address the reason for generation of supernumerary GFP foci in *cin8∆ kip3*∆ cells. At least two possibilities can be envisaged for this. Firstly, a leaky chromosome replication between meiosis I and II may amplify the operator arrays and cause >4 foci. However, this possibility seems unlikely because if there is a leaky replication of the operator array, due to close proximity (within 1.4 kb), the CenV would have been also replicated and in that case >4 kinetochore foci would have been observed which we failed to detect at any stage of meiosis (Fig. S3C). Secondly, due to an imbalance of spindle force acting on the centromeres, the chromosomes may break and since the operator arrays in our assays remain closed to the centromere or pericentromere, the arrays can also break to give >4 number of arrays and hence foci. Given the functions of the motors in moderating spindle-chromosome interaction through force generation, the latter possibility is more likely. To investigate if there is indeed any chromosome breakage, a single cell gel electrophoresis assay, known as comet assay, was performed (Ostling and Johanson, 1984). As it is difficult to lyse the tetrad because of the robust spore wall, cells were analyzed for the chromosome breakage at the tetra-nucleated stage before the formation of the spore wall. H_2_O_2_ (10mM) treated cells were used as a positive control for the breakage (Miloshev et al., 2002). Interestingly, we got a notable population of DNA masses that formed tails or comet phenotype in *cin8∆ kip3*∆ (approximately 20%) cells as compared to the wild-type (2.5%) or *cin8*∆ (1%) cells while in H_2_O_2_ treated sample, almost 46% of the cells exhibited the comet phenotype (Fig. 3A). These results suggest that the chromosome breakage does occur in *cin8∆ kip3*∆ cells.

**Figure 3:**
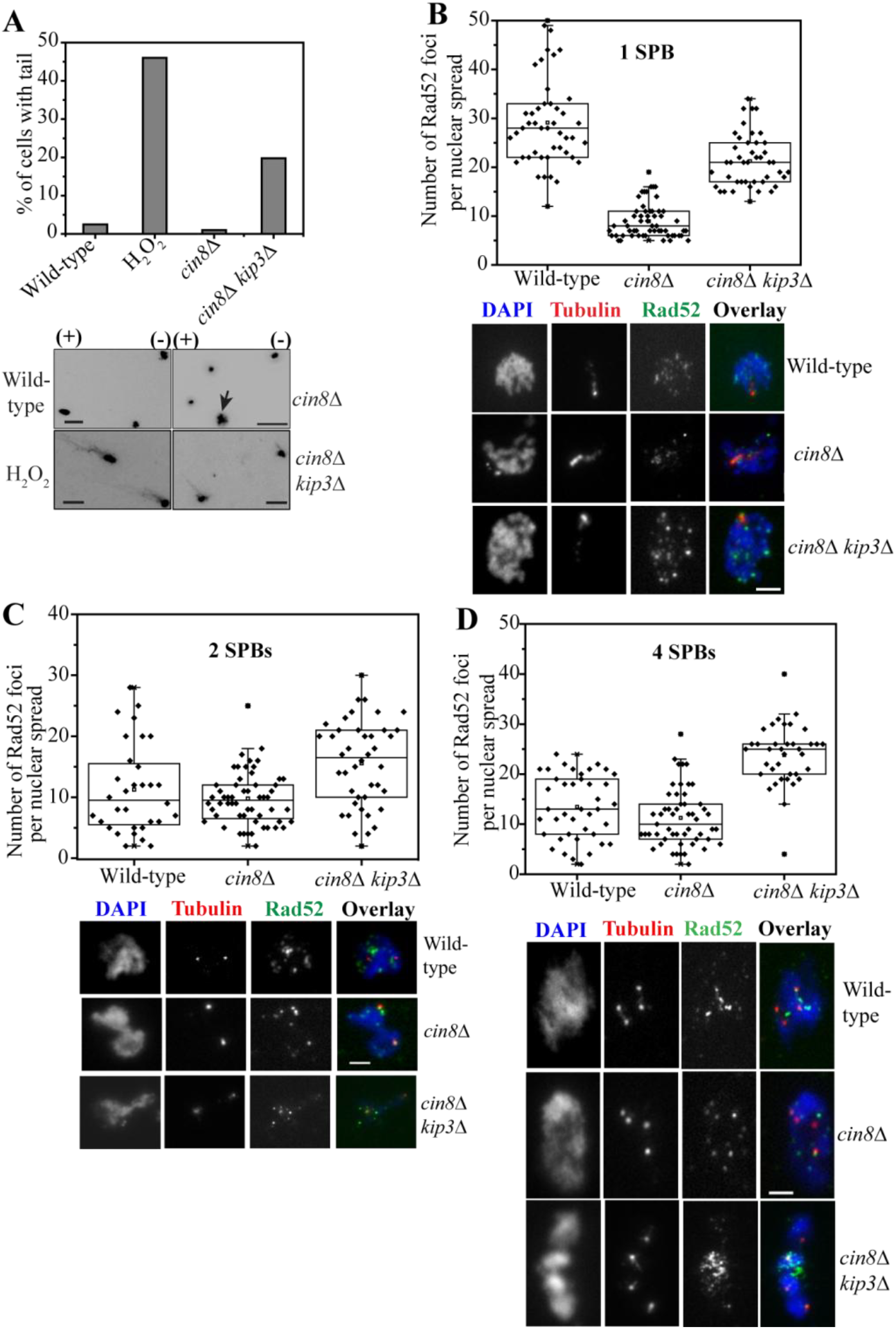
*cin8*∆ *kip3*∆ cells cause DNA breakage as they proceed through meiosis. (A) Meiotic cells were harvested for the comet assay (material and methods) from the SPM culture at the tetra-nuclated stage before the formation of the tetrads which is at 8 h for *cin8*∆ *kip3*∆ (SGY5089), at 7 h for *cin8*∆ (SGY315) and 5 h for wild-type (SGY40). For positive control, wild-type cells were treated with 10 mM H_2_O_2_ for 30 minutes at 4°C. The histograms correspond to the percentage of the cells that formed the comets. Representative gel images (with ‘+’ and ‘-’ polarities) show ethidium bromide stained DNA from each strain. Bar, 5 µm. Arrow indicates an unsphreoplasted cell. N = 80-156 nuclei were observed for the comet formation. (B, C, and D) Chromosome spreads show Rad52-EGFP foci stained using anti-GFP antibody in the wild-type (SGY5414), *cin8*∆ (SGY5422) and *cin8*∆ *kip3*∆ (SGY5415) strains. At least 40 spreads were analyzed for each type. Spindle poles marked by anti-α-tubulin antibody were used to judge the cell cycle stage of the spreads. Foci were counted after merging of the Z-stacks with the maximum intensities for tubulin and Rad52 keeping the threshold same for all the fields. 1 tubulin dot (1 SPB) within one DAPI mass represents the prophase I while 2 dots (2 SPBs, 1 dot each in two DAPI masses) represent anaphase I and 4 dots (4 SPBs, 1 dot each in four DAPI masses) represent meiosis II stages. Bar, 2 µm.

To further reconfirm the chromosome breakage, we looked at the localization pattern of Rad52 which is required for the repair of DNA double-stranded break (DSB) generated due to intrinsic or extrinsic factors (Barlow and Rothstein, 2009; Malone and Esposito, 1980; Orr-Weaver et al., 1981; Resnick and Martin, 1976). In meiosis, at prophase I, programmed DSBs occur for recombination and consequently Rad52 foci are visible at that stage in the wild-type. However, once the DSBs are repaired the average number of Rad52 foci reduces during later stages of meiosis (Gasior et al., 1998). We counted and compared the Rad52-EGFP foci in the wild-type, *cin8*∆ and *cin8*∆ *kip3*∆ chromosome spreads harboring 1 SPB within a single nucleus (prophase I stage, Fig. 3B), 2 SPBs, one in each of the 2 nuclei (anaphase I stage, Fig. 3C) and 4 SPBs, one in each of the 4 nuclei (anaphase II/post meiosis II stage, Fig. 3D). At prophase I, we observed an average 29 foci in the wild type which was reduced to 9 ± 3in *cin8*∆ spreads (Fig. 3B) which is consistent to the defective homolog pairing observed in *cin8*∆ (Fig. 1E). However, in *cin8*∆ *kip3*∆, the average count was nearly 21 which suggests that the loss of Kip3 by unknown mechanism rescues the defect of *cin8*∆. While analyzing the spreads at anaphase I, we noticed no significant difference in Rad52-EGFP staining between the wild-type and *cin8*∆ (wild-type 11±7; *cin8*∆ 10±4; Fig. 3C), but observed a slight increase in the staining in the double mutant (*cin8*∆ *kip3*∆ 15±7; Fig. 3C). However, a drastic accretion in Rad52-EGFP staining was observed in *cin8*∆ *kip3*∆ spreads at anaphase II/post meiosis II (tetra-nucleated stage) over the wild-type or the *cin8*∆ (wild-type 13±6, *cin8*∆ 11±6, *cin8*∆ *kip3*∆ 24±6; Fig. 3D). These results indicate that as the *cin8*∆ *kip3*∆ cells pass through meiosis II, they accumulate DNA damage in the form of DSBs and perhaps due to this the supernumerary CenV-GFP foci were observed only on meiosis II but not on meiosis I spindle (Fig. 2G).

Earlier we noticed supernumerary SPB formation in the kinetochore mutants as the cells enter into meiosis II (Agarwal et al., 2015). As both Cin8 and Kip3 also have some functional roles at the centromere, we reasoned that in *cin8*∆ *kip3*∆ cells, following interphase II (stage between meiosis I and II), may be >4 SPBs or spindle poles are generated and the resulting extra pole(s) may cause imbalance of force and hence chromosome breakage. However, analysis of the tetra-nulceated *cin8*∆ *kip3*∆ cells harboring >4 CenV-GFP foci showed only 4 SPBs (Fig. S3D) indicating at least multipolarity is not the cause of the chromosome breakage.

### *cin8∆ kip3*∆ hinders the cohesin removal from the chromatin in meiosis

In budding yeast cohesin is removed from the chromosome arms during anaphase I while the removal of the centromeric cohesin occurs during meiosis II. However, in the FEAR mutants the loss of cohesin from the arms delays as the meiotic cohesin protein, Rec8 was detected at the arm regions during anaphase I (Marston et al., 2003). Since we noticed *cin8*∆ *kip3*∆ exhibits the phenotypes similar to the FEAR mutants during meiosis (Fig. 1B, iii and 2B), we therefore, investigated if the double mutant is compromised in cohesin removal. We monitored the Rec8-EGFP staining at different stages of meiosis in the wild-type and *cin8*∆ *kip3*∆ cells. Meiotic stages were determined on the basis of number and distance between the Spc42 foci. The centromeric Rec8 was judged by its staining present only at the vicinity of the SPBs due to proximity of the centromeres to the SPBs whereas arm plus centromeric Rec8, termed as nuclear Rec8, was identified by its presence spanning a broader region between the two SPBs (Fig. 4A-B). We observed that in the wild-type, 64% of the anaphase I cells displayed centromeric Rec8 which was reduced to 35% in *cin8*∆ *kip3*∆ (Fig. 4C). However, we noticed more cells with nuclear Rec8 in the double mutant (65%) than the wild-type (36%) suggesting a defect is cohesin removal during metaphase I to anaphase I transition. Given cohesin removal completes during meiosis II, strikingly, nuclear Rec8 was observed even during meiosis II stage at a staggering population (45%) in *cin8*∆ *kip3*∆ whereas in the wild-type such population was insignificant (3%; Fig. 4A, B, and D). We obtained similar results in chromosome spreads immunostained for Rec8-EGFP where both the bi-nucleated and the tetra-nucleated spreads had high levels of nuclear Rec8 (82% and 54%, respectively) in *cin8*∆ *kip3*∆ with respect to the wild-type (19% and 11%, respectively; Fig. 4E-F). Notably, we observed that in *cin8*∆ also nuclear Rec8 persisted in a higher population of bi-nucleated spreads (70%; Fig. 4E). However, in majority of the spreads at the tetra-nucleated stage Rec8 appeared as a single dot indicating presence of negligible amount on the chromatin; in contrast a dispersed Rec8 signal all over the chromatin was observed in the double mutant in higher percentage of the spreads (Fig. 4F). These results altogether suggest that a prolonged cohesin-chromatin association throughout meiosis occurs in *cin8*∆, albeit at a lesser extent, and in *cin8*∆ *kip3*∆ cells. Due to this defect and the associated delay in spindle elongation and disassembly, *cin8*∆ and *cin8*∆ *kip3*∆ cells show a delay in meiosis I to meiosis II transition (Fig. 1B, iii). It is tempting to speculate that due to higher level of retention of cohesin in the *cin8*∆ *kip3*∆ cells, during anaphase I and anaphase II spindle elongations, the chromosomes cannot disjoin properly when subjected to pulling force exerted by the other motors and they eventually break causing very low spore viability.

**Figure 4:**
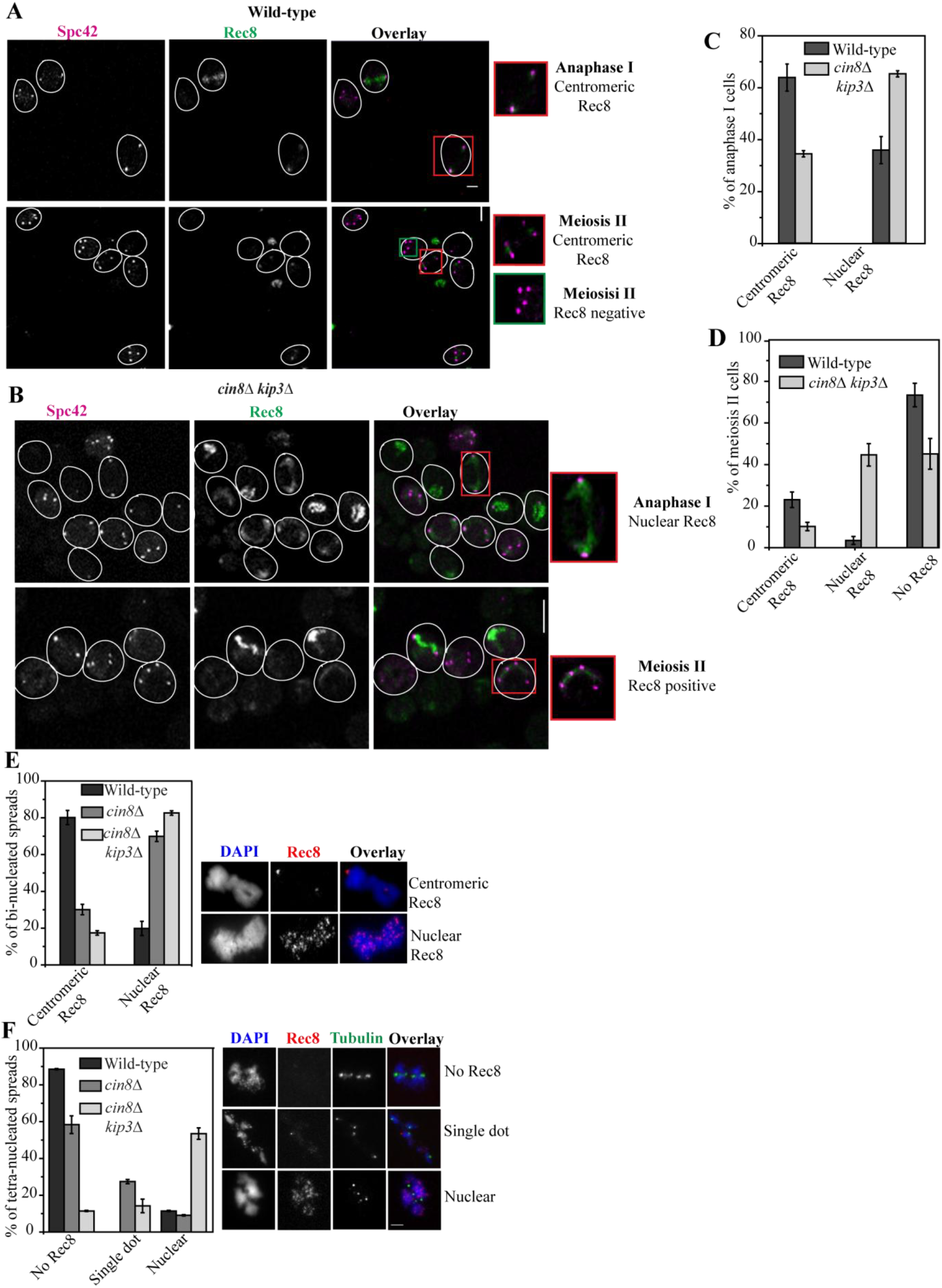
The removal of Rec8 cohesin is defective in *cin8*∆ *kip3*∆. (A – D) Wild-type (SGY5557) and *cin8*Δ *kip3*Δ (SGY5523) cells harboring Rec8-EGFP and Spc42-CFP were analyzed for Rec8 localization at different stages of meiosis. (A) The representative images where Rec8 staining appeared as two tight-knit dots at the vicinity of each of the two SPBs or as a single such dot each in between the two SPBs of two pairs were scored as centromeric Rec8 at anaphase I or meiosis II (metaphase II and anaphase II), respectively. Meiosis II cells with no Rec8 staining were also scored. (B) The representative images where Rec8 staining appeared in a broader region close to the SPBs or between the SPBS were scored as nuclear Rec8 (arm plus centromeric) at anaphase I or at meiosis II, respectively. (C - D) The percentages of anaphase I (C) and meiosis II (D) cells with the types of Rec8 staining are shown for the wild-type and *cin8*∆ *kip3***∆** cells. For (C) and (D) 148 and 267 cells were analyzed for the wild-type and *cin8*Δ *kip3*Δ, respectively. Bar, 5 µm. (E and F) Chromosome spreads from the wild-type (SGY5497; n = 83), *cin8*Δ (SGY5501; n = 114), and *cin8*Δ *kip3*Δ (SGY5500; n = 183) cells harboring Rec8-EGFP were monitored at different stages of meiosis by EGFP immunostaining (E) Quantitative analysis of Rec8 localization as only two foci (centromeric Rec8) or distributed throughout the chromatin (nuclear Rec8) in the bi-nucleated chromosome spreads. Representative image of each type is shown on the left. (F) Quantitative analysis of Rec8 localization as tiny (single dot) or large (nuclear) appearance on the chromosome spreads from the tetra-nucleates. Tetra-nucleated stage of the spreads was determined by tubulin immunostaining. *cin8*Δ *kip3*Δ cells show significant Rec8 throughout the chromatin (nuclear) even in the tetra-nucleated stage. Representative image of each type is shown on left. Bar, 2 µm.

As we observed a delay in spindle elongation and cell cycle progression in *cin8*∆ and *cin8*∆ *kip3*∆ cells, it is critical to address if the prolonged retention of Rec8 on the chromatin is due to a delay in degradation of securin (Pds1), a condition that releases separase to cleave Rec8. Under unperturbed condition Pds1 is degraded during metaphase I to anaphase I transition following re-appearance in metaphase II and degradation in anaphase II. We followed the level of Rec8 and Pds1 through synchronized meiosis in the wild-type, *cin8*∆, and *cin8*∆ *kip3*∆ cells through immunoblotting (Fig. 5A-B). As observed earlier (Fig. 1B) pace of meiosis was delayed in *cin8*∆ *kip3*∆ than the wild-type and *cin8*∆ (Fig. 5C), Pds1 degradation in the same strains also followed the same regime (Fig.5A-B). Notably, with disappearance of Pds1 all Rec8 was removed in the wild-type while the removal was deferred in *cin8*∆ and to a greater extent in *cin8*∆ *kip3*∆ (Fig. 5A-B). Consistent with our cell biological data (Fig. 4), we noticed that in the double mutant a significant Rec8 level was persistent even at 15 h in meiosis when around 90% of the cells had either entered into anaphase II or sporulated whereas in such cells either in the wild-type or in the *cin8*∆ cells, Rec8 was absent (Fig. 5A-C). Live cell imaging of Pds1-EGFP also revealed that there is no difference in Pds1 stability on anaphase I spindle between the wild-type and the double mutant (Fig. 5D). These results suggest that the protracted Rec8 retention on the chromatin in the *cin8*∆ *kip3*∆cells is not due to a biochemical delay imposed by persistent Pds1.

**Figure 5:**
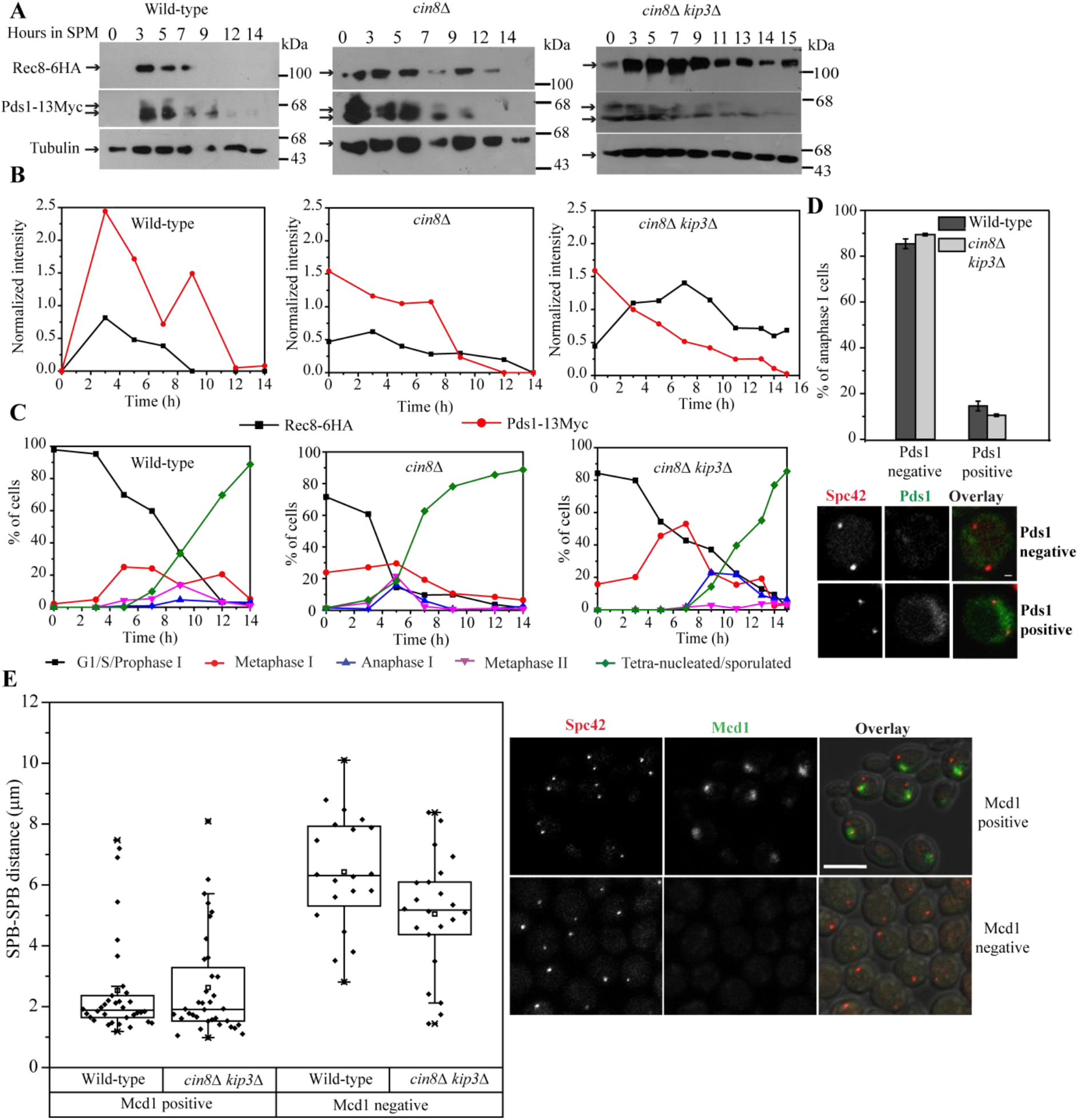
*cin8*∆ *kip3*∆ perturbs the co-ordination between Pds1 and Rec8 degradation in meiosis but does not affect the dynamics of Mcd1 removal in mitosis. (A) Western-blot analysis of the wild-type (SGY5534), *cin8*Δ (SGY5532), and *cin8*Δ *kip3*Δ (SGY5533) cells for the levels of Rec8-6HA and Pds1-13Myc at the indicated time points during meiotic progression. Tubulin was used as a loading control. (B) Densiometric analysis of Rec8-6HA and Pds1-13Myc signals obtained in (A) are shown after normalizing with the tubulin signal using ImageJ software. As evident from the graph, the drop in Rec8 level, unlike Pds1, is much slower in *cin8*Δ *kip3*Δ than in the wild-type or in *cin8*Δ. (C) The percentage of prophase I, metaphase I, anaphase I, metaphase II and anaphase II/sporulated cells were determined by tubulin immunostaining at the time points utilized for Rec8-6HA and Pds1-13Myc detection in (A and B). N ≥ 90 cells for each time point. (D) Analysis of Pds1-EGFP localization in the wild-type (SGY5570; n = 44) and *cin8*Δ *kip3*Δ (SGY5567; n = 75) cells during anaphase I stage determined by the distance between the two SPBs marked with Spc42-mcherry. (E) Localization of the Mcd1-EGFP in the wild-type (SGY5629) and *cin8*∆ *kip3*∆ (SGY5630) cells corresponding to the 3D distances between the two spindle poles marked by Spc42-mcherry. A field view of the metaphase (≤ 1.2 µm distance) and anaphase (≥ 2.5 µm distance) cells shows the presence and absence of Mcd1-EGFP, respectively. Error bars represent the standard deviation from the mean values obtained from three independent experiments. Bar, 2µm.

If the defect in cohesin removal in *cin8*∆ *kip3*∆ cells is responsible for chromosome mis-segregation, dyad formation and poor spore viability, the removal of Spo13, a meiosis specific protein that has a role in the centromeric cohesin protection during meiosis I (Shonn et al., 2002), in *cin8*∆ *kip3*∆ may improve the spore viability. Remarkably, the viability of the dyads was improved from 7% in *cin8*∆ *kip3*∆ to approximately 43% in *cin8*∆ *kip3*∆ *spo13*∆ (Fig. 2D). These data along with the cell biological and immunoblotting observations suggest that there is prolonged retention of Rec8 on the chromatin in the mutants and it is not due to cell cycle delay or delay in Pds1 degradation instead our results as described below indicate that perhaps some other factors also determine the fate of cohesin removal.

The failure in proper cohesin removal in post anaphase I *cin8*∆ *kip3*∆ cells instigated us to examine if the similar defect prevails in mitosis. For estimation of Mcd1 (mitotic cohesin) localization with respect to the distinct mitotic stages cells were released synchronously from G1 arrest and were examined for the presence or absence of Mcd1-GFP nuclear signal. With reference to the distances between the SPBs, we found no significant difference in the Mcd1-EGFP staining between the wild-type and *cin8*∆ *kip3*∆. The cells with the inter-polar distances in the range of 1.2 – 2.2 µm (metaphase/preanaphase) were positive for Mcd1-GFP while beyond that (post-anaphase) no Mcd1 staining was visible (Fig. 5E). These results indicate that mitotic cohesin removal is not perturbed in *cin8*∆ *kip3*∆ cells.

### Inhibition of recombination alleviates the anaphase I delay in *cin8*∆ *kip3*∆

The hindrance imposed by the reciprocal exchange (recombination) between the homologs restricting their disjunction is relieved by the dissolution of cohesin from the arm regions (Buonomo et al., 2000). Since, we observed a defect in cohesin removal and subsequent delay in anaphase I in the double mutant; removal of Spo11, the enzyme required for the double-stranded break formation to initiate recombination, in these cells should rescue the delay. We observed that the removal of Spo11 in the *cin8*∆ *kip3*∆ double mutant indeed caused the triple mutant to complete meiosis faster than the double mutant (Fig. S4A). For instance, at 5 h time point the percentage of anaphase I cells was reduced to approximately 4% in the triple mutant which was found 20% in the *cin8*∆ *kip3*∆ (Fig. S4A) suggesting a delay in the latter cells. Notably, there was a rise in the metaphase II population detected in the triple mutant (25%) over the other mutants that might be due to the combined effect of persistent cohesin and removal of Spo11 similar to what has been reported for the *P_CLB2_-CDC20 spo11*∆ cells (Lee and Amon, 2003). Nevertheless, these results further support that the cohesin dissolution is defective in *cin8*∆ *kip3*∆.

### *cin8*∆ *kip3*∆ causes homolog non-disjunction and aberrant meiosis II

As the defect in cohesin removal hinders homologue separation during meiosis I (Buonomo et al., 2000; Challa et al., 2019), we analyzed the homolog segregation in the bi-nucleated cells harboring homozygous CenV-GFP (Fig. 6A). Such cells with proper homolog disjunction will exhibit an equal number of CenV-GFP foci in each nucleus (2:2, Type I) whereas non-disjunction will result in unequal GFP foci distribution (1:0, 1:3, 4:0; Type II). We detected Type II phenotype in approximately 26% of *cin8*∆ *kip3*∆ cells over 12% in wild-type. Notably, a unique third category (around 17%, Type III) was observed only in *cin8*∆ *kip3*∆ where the CenV-GFP dots were present at the middle of a stretched DAPI. We believe the type III phenotype was generated as the sustained cohesin perturbs chiasmata resolution and impedes disjunction of the homolog ssince we observed a significant reduction in the distance between the two homologs in the bi-nucleated meiosis I cells (Figure 6B). In support of this, the removal of chiasmata by *spo11*∆ resulted in the reduction of the type III phenotype in *cin8*∆ *kip3*∆ *spo11*∆ cells (Fig. 6A). However, *spo11*∆ caused increased Type II frequency as loss of homolog pairing is known to perturb homolog bi-orientation and their disjunction (Buonomo et al., 2000; Klapholz et al., 1985).

**Figure 6:**
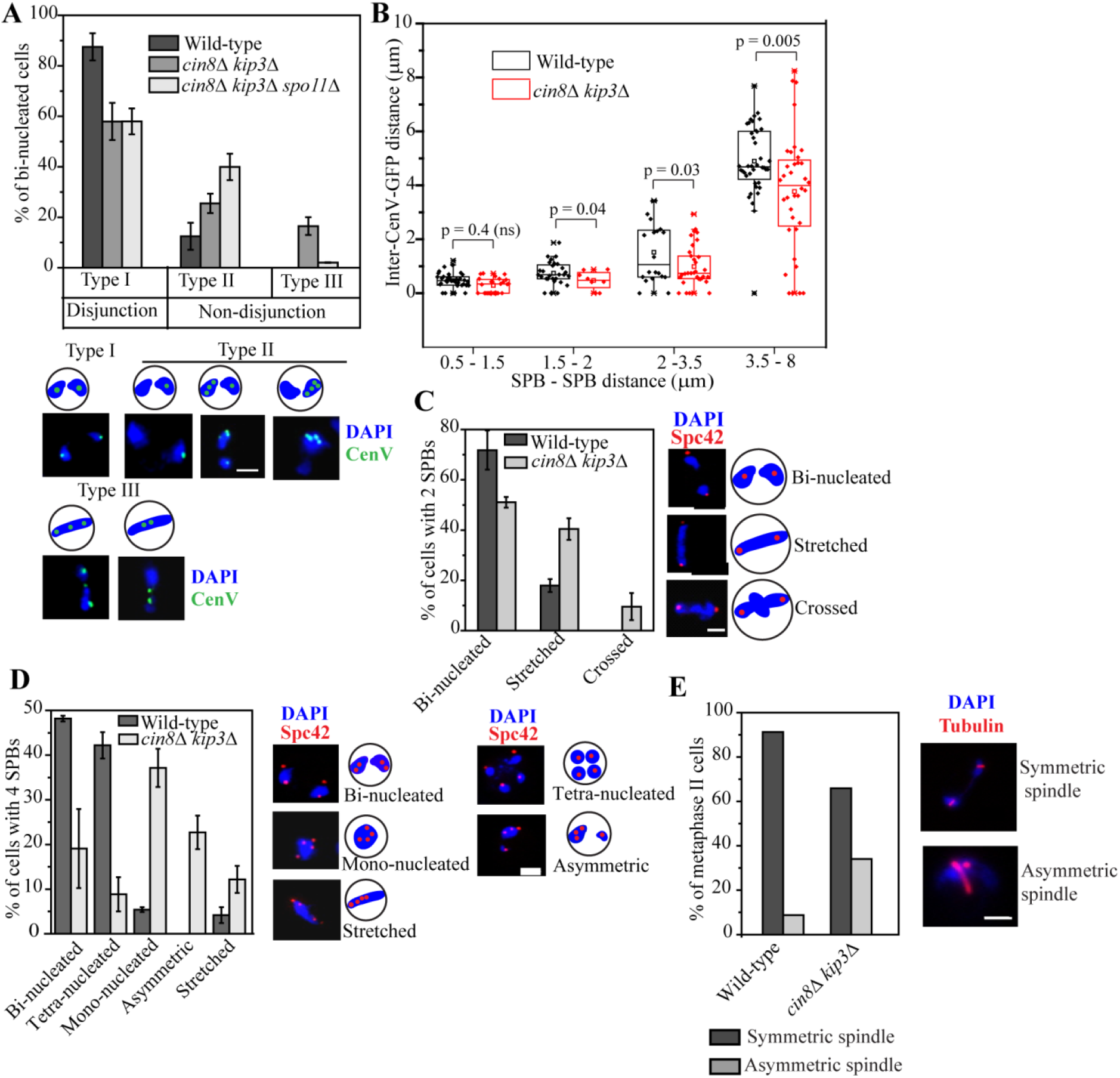
Homologous chromosomes and nuclear separation are impeded in *cin8*∆ *kip3*∆ cells. The wild-type (SGY9002) and *cin8*∆ *kip3*∆ (SGY5338) cells harboring homologously marked CenV-GFP and Spc42-mcherry were released into meiosis at 30° C and analyzed for the homolog (CenV) and nuclear segregation.(A) Segregation of the CenV-GFP homologs at the bi-nucleated stage are shown for the wild-type (n = 185), *cin8*∆ *kip3*∆ (n = 167) and *cin8*∆ *kip3*∆ *spo11*∆ (SGY5444; n = 97) cells. Representative images for the types of CenV homolog separation are shown. (B) The 3-D distances between the two CenV-GFP homologs are plotted with respect to the inter-polar (SPB-SPB) distances. The distances were measured using Imaris software (see materials and methods). Nuclear separation in the meiotic cells harboring 2 and 4 SPBs are depicted in (C) and (D), respectively (C, wild-type n = 70, *cin8*∆ *kip3*∆ n = 95; D, wild-type n = 166, *cin8*∆ *kip3*∆ n = 166). ‘n’ represents the total number of cells scored for the analysis. (E) Analysis of the tubulin morphology in the wild-type (SGY5001; n = 103) and *cin8*∆ *kip3*∆ (SGY5089; n = 91) meiosis II cells by tubulin immunostaining. Error bars represent the standard deviation from the mean values obtained from three independent experiments. p-value ≥ 0.05 is considered non-significant (ns). Bar, 2 µm.

A blockage in Rec8 cleavage in separase mutant, *esp1-1* hinders nuclear separation, however, following a prolonged arrest the cells embark on abrupt meiosis II (Buonomo et al., 2000). Since homolog non-disjunction was found impaired in *cin8*∆ *kip3*∆ bi-nucleated cells which include both anaphase I as well as metaphase II cells (Fig. 6A), we examined meiosis I and II nuclear segregations in the cells harboring two (Fig.6C) and four (Fig. 6D) SPBs, respectively. Given that cohesin retention in *cin8*∆ *kip3*∆ is not due to Pds1 stability (Fig. 5A-D), we argued that these cells would progress through meiosis I in spite of having physical barrier in nuclear separation. As expected we observed a significant population of the post-anaphase I cells in *cin8*∆ *kip3*∆ within complete nuclear division as evident from the ‘stretched’ nuclear morphology (approximately 40%; Fig. 6C).

We also observed a meager population (approximately 10%) of anaphase I cells with three connecting nuclear lobes (‘crossed’ morphology) only in *cin8*∆ *kip3*∆. This category of DAPI segregation resembles the one obtained in FEAR mutants resulting from the initiation of meiosis II on the meiosis I spindle (Marston et al., 2003). The population of cells under ‘stretched’ and ‘crossed’ categories either evade meiosis II forming dyads (Fig. 2B) or they abruptly enter into meiosis II where they mostly showed asymmetric (26%) and no nuclear separation (mono-nucleates; 41%) with 4 SPBs (Fig.6D). Similar phenotypes were observed in *mam1*∆ cells due to delayed nuclear division (Toth et al., 2000). Further, due to prolonged anaphase I and subsequent abrupt initiation of meiosis II, there was a significant difference in the length of the two spindles in meiosis II in around 34% *cin8∆ kip3*∆ cells (Fig. 6E). This phenotype is similar to the one observed in meiosis II cells of *mam1*∆ in budding yeast (Toth et al., 2000) and recombination-defective mutant, *rec8* in *S. pombe* (Yamamoto et al., 2008) where the common responsible factor is the delayed nuclear separation.

### Kip1 degradation is delayed in *cin8*∆ *kip3*∆

In *cin8*∆ *kip3*∆ cells, although delayed, spindle elongation does occur and we believe that Kip1 executes this function in a protracted way. In mitosis, Kip1 is degraded during the onset of anaphase by Cdc20 (Hildebrandt and Hoyt, 2001). To investigate if Kip1 becomes more stable in the absence of Cin8 and Kip3, we compared the Kip1 level by immunoblotting between the wild-type and *cin8*∆ *kip3*∆ cells during different stages of meiosis (Fig.7A-B). Given the difference in the pace of the cell cycle, the 10 h stage of wild-type was considered equivalent to the 12 h of *cin8*∆ *kip3*∆ as the percentage of tetra-nucleated cells were observed almost similar (Fig. 7C). Kip1 was found stable for longer duration in *cin8*∆ *kip3*∆ than in the wild-type (Fig. 7A-B). To further examine this we monitored the localization of Kip1 in the wild-type and *cin8*∆ *kip3*∆ undergoing meiosis using live cell imaging. Stages were judged on the basis of number of SPB and the distance between two SPBs’. In anaphase I cells, Kip1 was either localized along the spindle (42%) or near the poles (12%) while in 46% of cells Kip1 was absent suggesting it degrades towards the end of meiosis I. In contrast, Kip1 was absent in only 4% of anaphase I cells in *cin8*∆ *kip3*∆ (Fig.7D). In metaphase II while almost 100% wild-type cells showed polar localization of Kip1, almost 45% of the *cin8*∆ *kip3*∆ cells exhibited single spindle-like localization spanning the 4 SPBs suggesting that Kip1 degradation is defered in the latter cells (Fig. 7E). However, overall Kip1 level was not altered as determined by comparing Kip1-EGFP intensity between the wild-type and *cin8*∆ *kip3*∆ at anaphase I and metaphase II (Fig. 7F-G). For intensity measurement within anaphase I cells only spindle-localized Kip1-EGP intensity was compared between the wild-type and the mutant. These results indicate that in absence of Cin8 and Kip3, spindle elongation can still be possible perhaps through positive regulation of Kip1 function. The presence of Kip1 along the spindle in metaphase II *cin8*∆ *kip3*∆ cells points towards longer stability of this protein.

**Figure 7:**
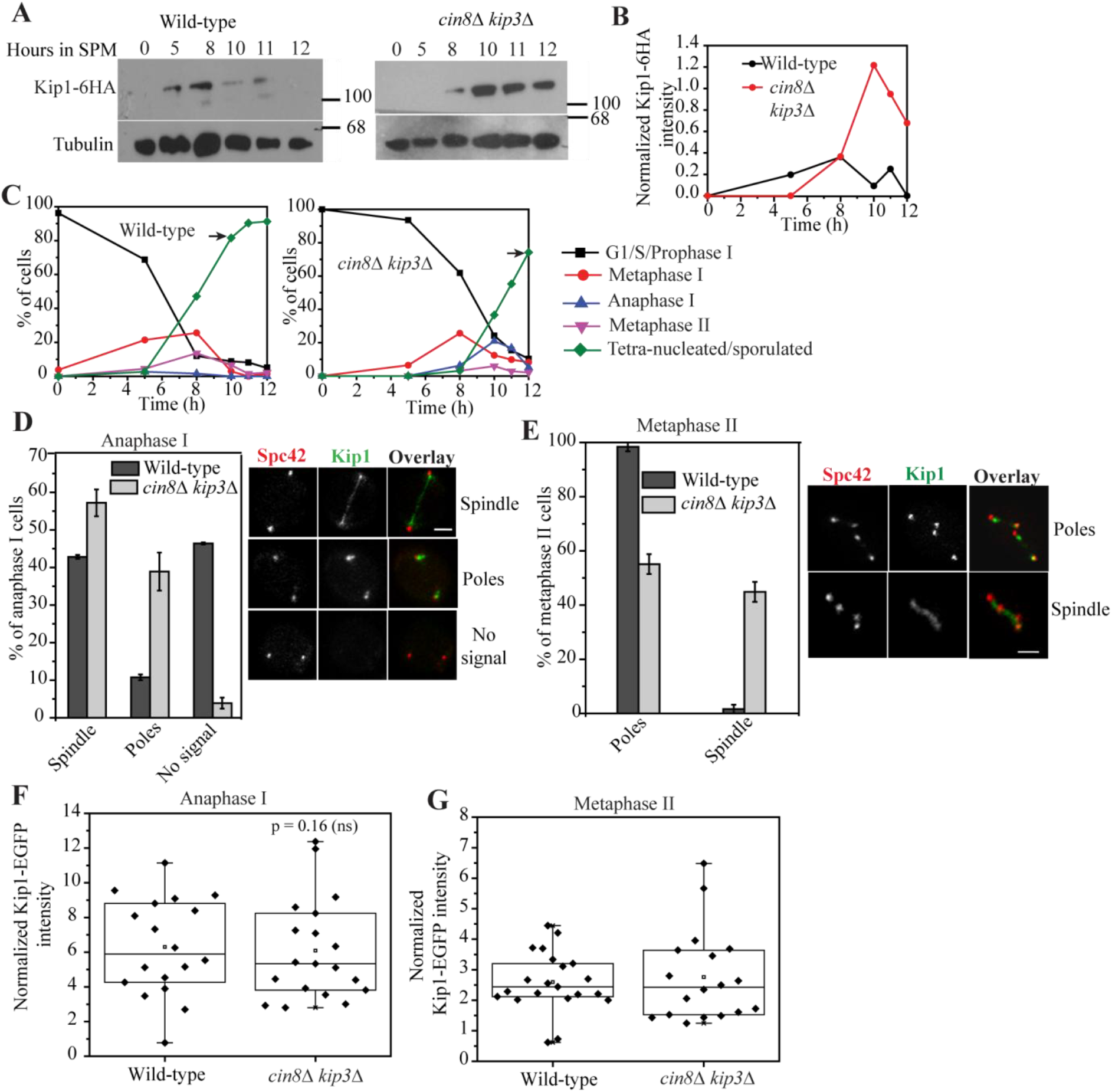
Increased stability of Kip1 in the *cin8*∆ *kip3*∆ cells during meiosis. (A) The **w**ild-type (SGY5539) and *cin8*∆ *kip3*∆ (SGY5540) cells harboring Kip1-6HA were induced for synchronized meiosis and analyzed for the levels of Kip1-6HA at the indicated time points during meiotic progression. Tubulin was used as a loading control. (B) Densiometric analysis of Kip1-6HA bands obtained in (A) after normalization with the respective tubulin bands using ImageJ software. (C) The percentages of prophase I, metaphase I, anaphase I, metaphase II and anaphase II/sporulated cells were determined by tubulin immunostaining in the wild-type and *cin8*∆ *kip3*∆ cells at the time points utilized for Kip1-6HA detection in (A and B). ‘n’ ≥90 cells for each time point. Arrow indicates a reference time-point for comparison between the wild-type and *cin8*∆ *kip3*∆. The localization of Kip1-EGFP in the wild-type (SGY5051) and *cin8*∆ *kip3*∆ (SGY5561) cells is shown during anaphase I (D; wild-type n = 56, *cin8*∆ *kip3*∆ n = 97) and metaphase II (E; wild-type n = 47, *cin8*∆ *kip3*∆ n = 165). The stage of a meiotic cell was determined by the number and distance between the SPBs. ‘n’ represents the number of cells analyzed for the assay. The dot plots of Kip1-EGFP intensities in the wild-type and *cin8*∆ *kip3*∆ cells are shown during anaphase I (F) and metaphase II (G). Each signal intensity value was normalized with the background and Spc42-mcherry intensity values. Error bars represent the standard deviation from the mean values obtained from three independent experiments. p-value ≥ 0.05 was considered non-significant (ns). Bar, 2 µm.

### The tension generated by the microtubule mediated force drives efficient Rec8 removal

From the above results, it is evident that in the absence of both Cin8 and Kip3, Rec8 is not efficiently removed from the chromatin and that condition perhaps leads to chromosome breakage during meiosis II. What could be the reason for Rec8 retention when Cin8 and Kip3 are not present? We argue that in *cin8*∆ *kip3*∆ cells due to the absence of microtubule cross-linking and depolymerization activities, there is inadequate microtubule based pulling force acting on the kinetochores and consequently the chromatids and the cohesin between the sisters are not under sufficient tension both in meiosis I and meiosis II. Given this, we hypothesize that generation of tension on cohesion is perhaps a novel determinant for efficient Rec8 removal. If this is true, then the generation of microtubule force in the *cin8*∆ *kip3*∆ cells can rescue the Rec8 cleavage and therefore chromosome integrity and spore viability. To test this, we expressed in these cells a phosphodeficient allele of *CIN8* (Cin8-3A) that retains on the spindle and can generate force for an extended period or a phosphomimic allele of *CIN8* (Cin8-3D) that fails to bind to the microtubule and create force and thus exhibits a diffused nuclear localization (Avunie-Masala et al., 2011). The spindle localization of Cin8-3A while a diffused localization of Cin8-3D in the anaphase I cells confirmed their modes of action (Fig. 8A). Remarkably, in the chromosome segregation assay with homozygous CenV-GFP, we observed a drop in the percentage of tetra-nucleates harboring >4 GFP dots, a readout of chromosome breakage, in *cin8*∆ *kip3*∆ Cin8-3A cells (15%) compared to *cin8*∆ *kip3*∆ (29%; Type IV, Fig.8B). In accord to this the spore viability obtained in *cin8*∆ *kip3*∆ (approximately 16%) was ameliorated to a great extent upon expression of Cin8-3A (approximately 48%, Fig. 8C). The observed rescue effect is specific to the ability of Cin8-3A to bind to the microtubule as *cin8*∆ *kip3*∆ cells expressing Cin8-3D showed similar phenotypes to that of *cin8*∆ *kip3*∆ alone (Fig. 8B-C). These results indicate that the tension generated by Cin8 and Kip3 collectively via microtubule perhaps create a signal for efficient cleavage and subsequent removal of Rec8.

**Figure 8:**
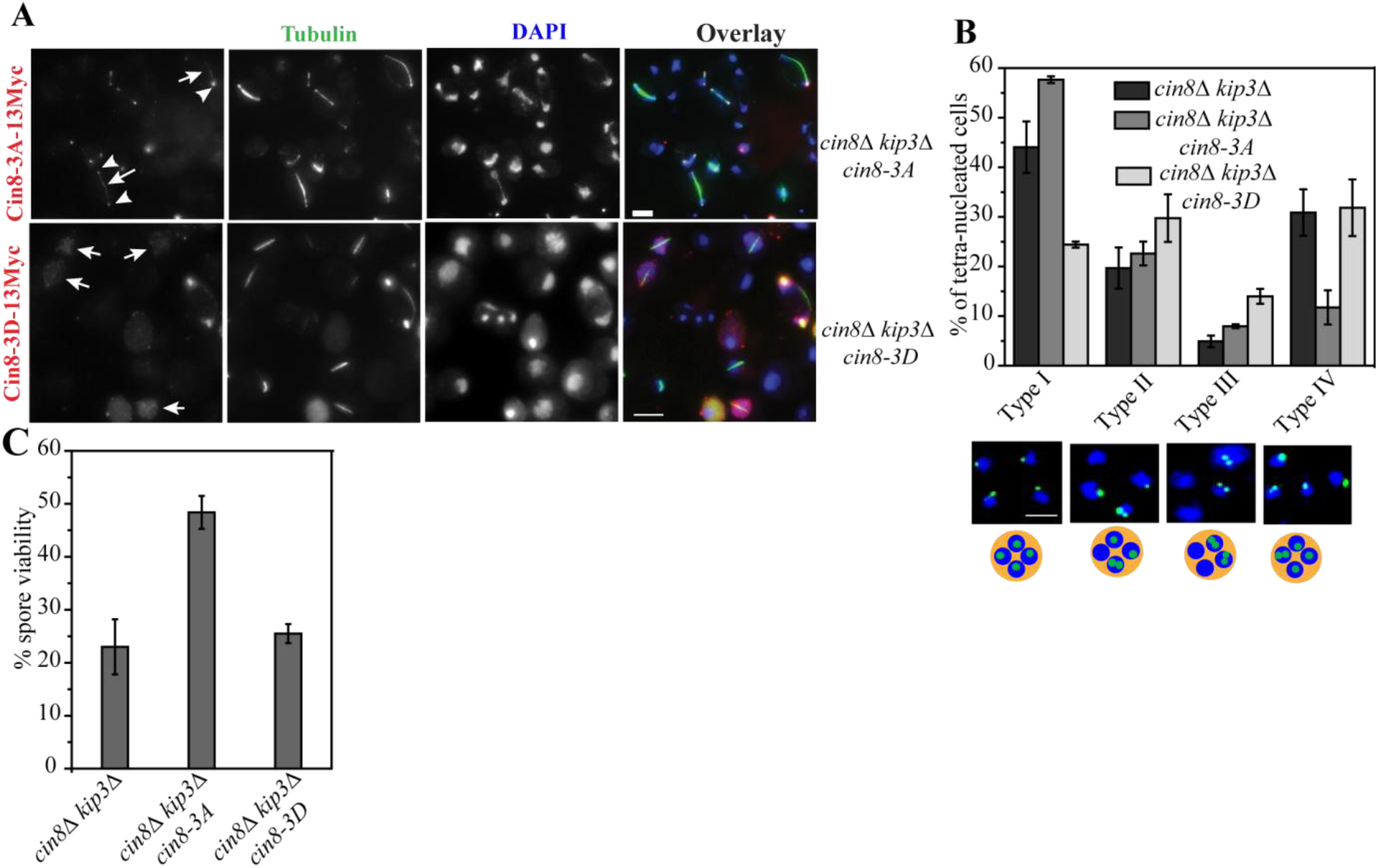
The absence of microtubule cross-linking activity of Cin8 is partly responsible for the defect observed in *cin8*∆ *kip3*∆. **(A)***cin8*∆ *kip3*∆ *cin8-3A-13MYC* (SGY5546), and *cin8*∆ *kip3*∆ *cin8-3D-13MYC* (SGY5547) cells expressing phosphodeficient or phosphomimic alleles of *CIN8*, respectively were examined during meiosis for the localization of the corresponding mutant proteins with respect to the spindle. While Cin8-3A-13Myc was detected along the long spindles (white arrows) and at the poles (white arrowheads), Cin8-3D-13Myc was observed as diffused nuclear signals (red arrows). Bar, 5 µm. (B) Homologous CenV-GFP segregation in the tetra-nucleated cells of the above two strains and *cin8*∆ *kip3*∆ (SGY5089) strain are shown. More than 80 tetra-nucletes were counted for each strain. The frequency of tetra-nucleates with supernumerary GFP dots (Type IV) was reduced in the cells expressing Cin8-3A that can bind and cross-link the microtubules. Bar, 2 µm (C) The percentage of spore viability in the above strains. More than 70 tetrads were dissected for each strain. Error bars represent the standard deviation from the mean values obtained from three independent experiments. Bar, 2 µm.

To further test whether tension is an additional factor required for the cohesin removal in meiosis, we monitored the Rec8 localization after mimicking the loss of tension condition by two distinct approaches. During meiosis I, the tension between the homologs and on the cohesin is generated as the bipolar pulling force by the microtubule is opposed by chiasmata formed between the non-sister chromatids and the cohesion formed between the sisters and the non-sisters. We inhibited the chiasmata formation by deleting *SPO11* and examined the Rec8 localization and compared that with the previous results (figs. 9A and 4A-D). Nuclear Rec8 localization was observed in around 92% of the *spo11*∆ anaphase I cells, which was far more than observed in *cin8*∆ *kip3*∆ (65%) or in wild-type (36%; figs. 9A and 4A-C) suggesting loss of tension indeed resists efficient cohesin removal. However, as the *spo11*∆ cells proceeded to meiosis II, Rec8 staining pattern in metaphase II became similar to the wild-type cells. This was expected since *spo11*∆ can alleviate tension only during meiosis I. In another approach to investigate the role of tension in Rec8 removal, we depolymerized the microtubules using benomyl (materials and methods) in the cells depleted for spindle assembly checkpoint protein Mad2 using *CLB2* promoter (Jin et al., 2009) so that the cells can proceed through meiosis. We treated the cells with benomyl after 5.5 h of meiotic release when most of the cells have passed the prophase I stage (Fig. 9B). In absence of Mad2, benomyl treated cells were able to go through meiosis I and meiosis II although not as efficient as the mock-treated cells (Fig. 9C). Due to absence of microtubules as the SPB separation was improper, we were unable to distinguish between the metaphase I and anaphase I cells and therefore only the cells with 4 SPBs were analyzed. We observed that a notable population (69%) of cells harbored robust nuclear Rec8 staining in the benomyl treated culture but no or minimal centromeric Rec8 staining in the mock-treated culture (Fig.9D). This suggests that the removal of microtubules by benomyl reduces tension and that in turn perturbs Rec8 cleavage. Consequently, it is expected that the benomyl treated cells harboring homozygous CenV-GFP would cause chromosome breakage during meiosis II and show supernumerary GFP foci. Although the DAPI segregation in the presence of a sublethal concentration of benomyl was not as efficient as the mock treated culture, we observed around 32% of tetra-nucleated cells with supernumerary GFP foci in the presence of the drug which was meager 5% under unperturbed condition (Fig. 9E). The above two investigations indicate that the reduction of tension can cause inefficient cohesin removal and we suggest that this condition eventually leads to chromosome breakage as observed in the *cin8*∆ *kip3*∆ cells.

**Figure 9:**
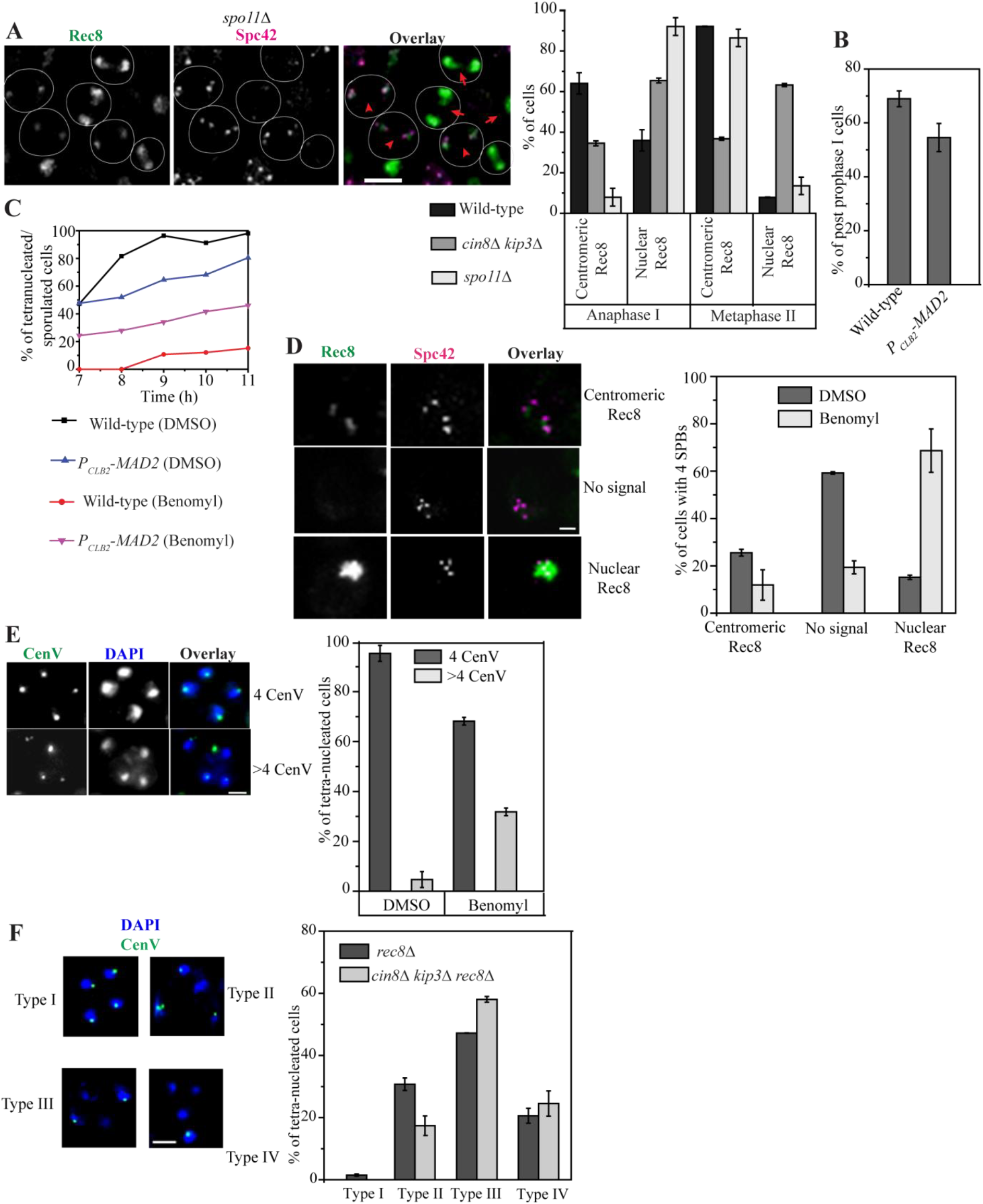
Tension is indispensable for the timely removal of Rec8 and deletion of *REC8* suppresses formation of the supernumerary centromeric foci in *cin8*∆ *kip3*∆ during meiosis. (A) The localization of Rec8-EGFP in *spo11*∆ (SGY5610; n = 220) harboring Spc42-CFP in anaphase I and metaphase II stages with respect to the wild-type and *cin8*∆ *kip3*∆ as shown in figure 4C-D. The representative images for *spo11*∆ are shown on left whereas the same for the wild-type and *cin8*∆ *kip3*∆ are shown in figure 4A and 4B, respectively. Based on the distribution of Rec8 staining, cells were categorized as centromeric or nuclear Rec8 as shown by red arrowheads or arrows, respectively in the representative images. (B-C) Mad2 depletion relieves the cell cycle arrest caused by microtubule disruption. The wild-type (SGY5557) and *P_CLB2_-MAD2* (SGY5628) cells harboring Rec8-EGFP and Spc42-CFP were released into synchronized meiosis. The progression of the cells through meiosis before or after the addition of benomyl was analysed by tubulin immunostaining and DAPI staining. (B) Following 5.5 h of meiotic release into the drug free medium, the percentages of cells that have progressed beyond prophase I are shown. (C) Each meiotic culture was then either treated with mock (DMSO) or benomyl and percentages of tetra-nucleated/sporulated cells at the indicated time points are shown. (D) Localization of Rec8-EGFP in the Mad2 depleted cells harboring *P_CLB2_-MAD2* (SGY5628) in the presence or absence of benomyl. Cells showing 4 SPBs marked by Spc42-CFP were scored for centromeric or nuclear or no signal of Rec8. (E) *P_CLB2_-MAD2* cells harboring homozygous CenV-GFP (SGY3248) were analyzed for the segregation of CenV-GFP at the tetra-nucleated stage following microtubule depolymerization by benomyl. For (D) and (E), around 120 - 150 cells were analyzed from the benomyl treated or untreated culture and the drug/DMSO was added following 5.5 h of meiotic induction. (F) Analysis of the CenV-GFP marked homolog segregation in the tetra-nucleates of *rec8*∆ (SGY5667; n = 142) and *cin8*Δ *kip3*Δ *rec8*∆ (SGY5670; n = 74) cells. ‘n’ represents the total number of tetra-nucleates scored for chromosome segregation. Error bars represent the standard deviation from the mean values obtained from three independent experiments. Bar, 2 µm.

If retention of Rec8 is responsible for chromosome breakage in the *cin8*∆ *kip3*∆ cells, then removal of Rec8 in these cells should alleviate the defect. To examine this, we deleted *REC8* in the wild-type and *cin8*∆ *kip3*∆ strains harboring homozygous CenV-GFP. Since meiosis is severely compromised in the absence of Rec8 (Klein et al., 1999), we observed very less population of tetra-nucleates in *rec8*∆ or in *cin8*∆ *kip3*∆ *rec8*∆. Due to high rate of chromosome non-disjunction in absence of cohesin, the percentage of tetra-nucleates with GFP dots in all the four nuclei was negligible (approximately 1%, Type I, Fig. 9F); instead we observed a predominant population of tetra-nucleates with GFP dots in 2 nuclei in *rec8*∆ and *cin8*∆ *kip3*∆ *rec8*∆ cells (47% and 58%, respectively, Type III) while the remaining population contained GFP dots either in three of the four nuclei (31% and 17% in *rec8*∆ and *cin8*∆ *kip3*∆ *rec8*∆, respectively, Type II) or only in one of the four nuclei (21%and 25% in *rec8*∆ and *cin8*∆ *kip3*∆ *rec8*∆, respectively, Type IV). This gross chromosome mis-segregation was also evident from the asymmetric DAPI staining observed in the tetra-nucleates. However, as we expected, none of the triple mutant cells exhibited >4 CenV-GFP dots indicating that the defective cohesin removal in the *cin8*∆ *kip3*∆ cells is indeed responsible for the chromosome breakage which is also depicted in our model (Fig. 10).

**Figure 10:**
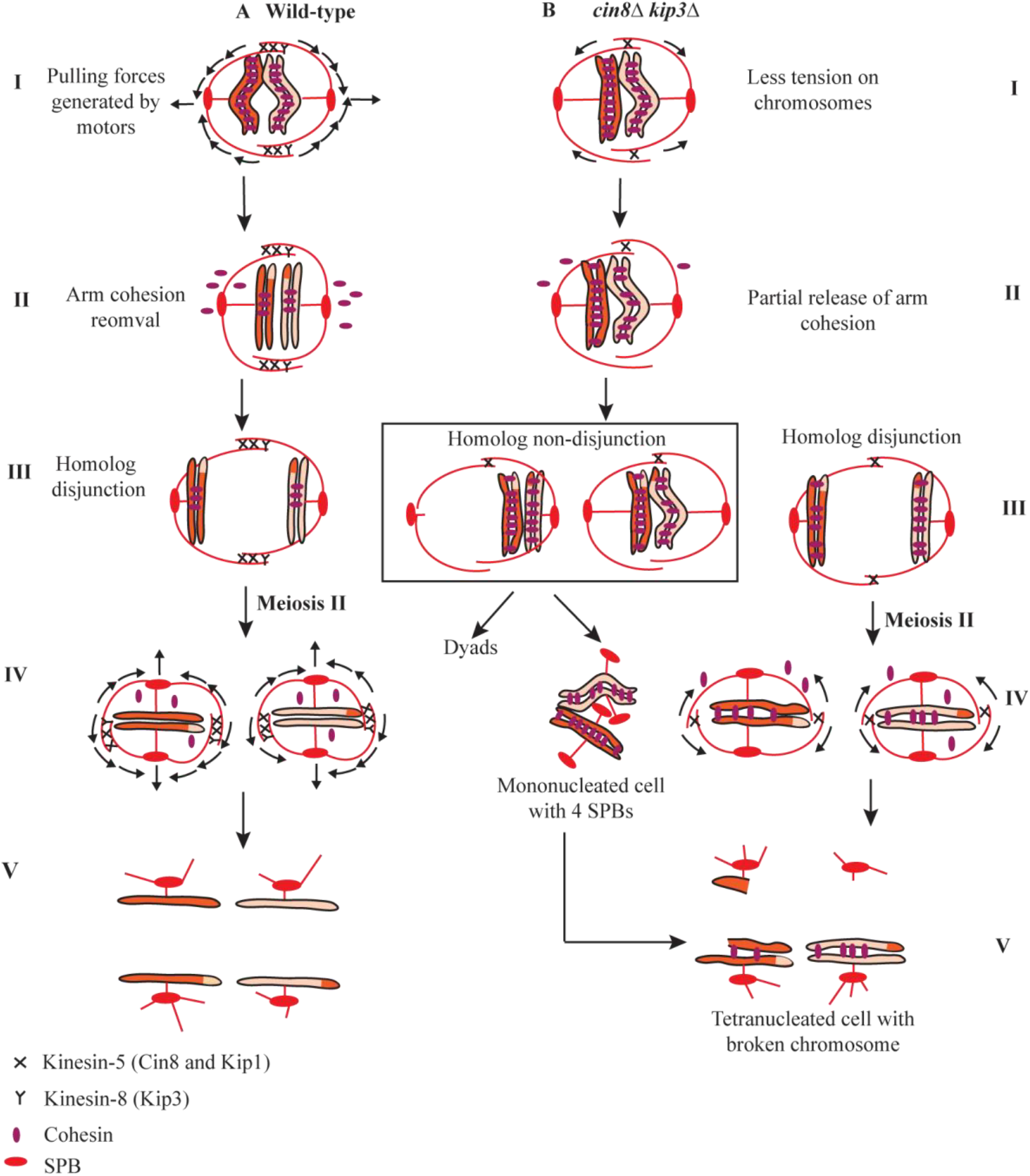
A possible mechanism responsible for the chromosome breakage in *cin8∆ kip3*∆. Model showing the meiotic chromosome segregation in the (A) wild-type and (B) *cin8∆ kip3∆.* (I) Due to absence of Cin8 and Kip3, the transduction of pulling force on the chromosomes by only Kip1 through sliding of the anti-parallel microtubules is less in the double mutant as compared to the wild-type. (II) This causes a lack of tension on the cohesins perturbing their removal. (III) The persistence of cohesin resists chiasmata resolution which along with weak kinetochore-microtubule attachment in *cin8∆ kip3*∆ results in homolog non-disjunction These cells after a transient delay at anaphase I either form dyads or enter into meiosis II showing mono-nucleates with 4 SPBs perhaps due to structural blockage in disjoining chromatids which can subsequently produce tetra-nucleates with broken chromosomes. However, in some cells homolog disjunction occurs but with persistent cohesin (IV) which results in chromosome breakage when the cells enter into meiosis II (V).

Shugoshin, Sgo1 is known to promote pericentromeric cohesin protection by reversing Rec8 phosphorylation through recruitment of PP2A (protein phosphatase 2A) at the centromere (Riedel et al., 2006). Sgo1 localization at the pericentromere is tension dependent and its dislodgement from there occurs when the kinetochores are under tension (Nerusheva et al., 2014). Since we observed that the absence of tension causes Rec8 protection in the *cin8*∆ *kip3*∆ cells, we argued that in these cells Sgo1 may be excessively associated with the chromatin driving the protection. However, we observed no significant difference in the Sgo1-6HA staining profile in the chromatin spreads between the wild-type and *cin8*∆ *kip3*∆ cells at the bi-nucleated or the meiosis II (post-anaphase I) stage (Fig. S4B-C). Sgo1 localized predominantly as clustered foci near the spindle poles in the anaphase I (Type I, Fig. S4B), and meiosis II spreads (Type I, Fig. S4C) while in the remaining population multiple foci of Sgo1 (Type II) were visible which presumably represented the pericentromeric distribution of Sgo1 as reported earlier (Katis et al., 2004).

## Discussion

Faithful chromosome segregation relies on a coordinated interaction between the microtubule and the chromosomes. The molecular motors profoundly influence this interaction not only by ensuring proper attachment of the kinetochores to the microtubules but also by temporally and spatially regulating the microtubule spindle. Budding yeast has four nuclear kinesin motors whose functions in the context of mitotic chromosome segregation have been described in several studies (DeZwaan et al., 1997; Hildebrandt and Hoyt, 2000; Roof et al., 1992; Winey and Bloom, 2012). However, given the differences in the pattern of chromosome movement between mitosis and meiosis including two-time chromosome segregation with concomitant assembly, extension and disassembly of the spindle in meiosis, it is intriguing to investigate the functions of these motors in meiosis. In this work, we analyzed the functions of the three microtubule plus end directed kinesin motors in meiosis.

### Loss of Cin8 perturbs homolog pairing and homolog disjunction

Analysis of the single motor mutants revealed that the loss of Cin8 affects meiosis more than the loss of either Kip1 or Kip3 (Fig. 1). At early meiosis, Cin8 appears to promote homolog pairing and consequently homolog disjunction (Fig. 1E-G) without any apparent role in sister chromatid cohesion or their mono-orientation (Fig. S1). However, it is possible that the roles of the motors in these events are not adequately unmasked due to functional redundancy among these motors. In homolog pairing, it is required that each homolog should locate each other which requires the functions of the cytoskeleton and the motors that are believed to facilitate pairing by enhancing the search rate. The function of the dynein motors with the help of nuclear envelope spanning SUN/KASH proteins in homolog pairing has been demonstrated in *C. elegans* (Penkner et al., 2009; Sato et al., 2009). In *S. pombe*, the dynein motors drive the ‘horsetail nuclear movement’ that facilitates homolog pairing (Wells et al., 2006). In *S. cerevisiae*, the rapid prophase movements (RPMs) of chromosomes in meiosis I is believed to occur via actin and nuclear envelope motor proteins including Mps3-Ndj1-Csm4 through interactions with the telomeres (Conrad et al., 2008; Koszul et al., 2008; Wanat et al., 2008). While RPM and the telomere-led movements of the chromosomes promote homolog pairing, it is plausible that the nuclear kinesins may facilitate RPM or they may function in pairing independently. However, the former possibility is unlikely since no interaction among the nuclear envelope proteins and the kinesin motors has been demonstrated. With a better microtubule cross-linking activity than the kinesin-8 motor (Kip3), the kinesin-5 motors (Cin8/Kip1) may have more role in the movement of one homolog with respect to the other during the search for the pairing partner. The fact that we observed a more significant effect of Cin8 over Kip1 in homolog pairing and for that matter in meiosis (Fig. 1E) despite both are of kinesin 5 family, is not surprising as in mitosis it has been demonstrated that Cin8 plays a larger role than Kip1 in chromosome segregation (Geiser et al., 1997) which is perhaps due to structural difference between these two proteins (Gerson-Gurwitz et al., 2011; Singh et al., 2018).

### Cin8 and Kip3 together are essential for timely exit from meiosis I and completion of meiosis II

It is intriguing that a gross drop in spore viability occurs in *cin8*∆ *kip3*∆ but not in *kip1*∆ *kip3*∆ double mutant (Fig.1A). Further analysis revealed that the *cin8*∆ *kip3*∆ cells share the phenotypes of the FEAR mutants that include a delay in spindle elongation and disassembly and generation of dyads (Figs. 1, ii-iii and 2B). We believe that this happens because Cin8 dephosphorylation by the FEAR-released Cdc14 is essential for maintaining Cin8 at the spindle (Roccuzzo et al., 2015). Due to lack of Cdc14 in the FEAR mutant, the phosphorylated form of Cin8 by Cdk1 is enriched which dissociates Cin8 from the spindle (Avunie-Masala et al., 2011). Therefore, the spindle phenotypes observed in the FEAR mutants of the meiotic cells resemble to the cells devoid of both Cin8 and Kip3. However, notably *cin8*∆ mutant alone does not show a FEAR-like phenotype indicating that Kip3 function is parallel to Cin8 at least at the spindle and its function might be similarly modulated by the absence of Cdc14. Although Kip3 function has not been reported to be regulated by Cdc14, in a screen using yeast proteomic library, Kip3 was identified as one of the Cdk1 substrates (Ubersax et al., 2003). Given Cdc14 is known to undo most of the Cdk1 mediated phosphorylations and in *S. pombe*, one of the kinesin-8, Klp-6 is a substrate of Cdc14 homolog Clp1 (Chen et al., 2013), it is possible that Cdc14 might regulate the Kip3 function in *S. cerevisiae*. In addition, similar to the FEAR mutants, *cin8*∆ *kip3*∆ cells mostly showed reductional segregation in the two spores of the dyads. However, some cells did complete meiosis II and produce tetra-nucleates but with dire consequences as discussed below.

### Improper cohesin removal in *cin8∆ kip3*∆ cells causes chromosome breakage in meiosis

The finding of >4 CenV-GFP foci in the homozygously GFP marked *cin8*∆ *kip3*∆ cells specifically in meiosis but not in mitosis was surprising (Figs. 2E-F and Fig. S3B). Further analysis revealed that this happens due to chromosome breakage in those cells that attempt to complete meiosis II (Fig. 2G). Unexpectedly, our investigations suggest that this breakage is due to improper removal of cohesin from the chromatin during both metaphase I to anaphase I and metaphase II to anaphase II transitions (Figs. 4 and 5A-D). We believe that anaphase I delay in *cin8*∆ *kip3*∆ cells, besides lack of sliding of the antiparallel microtubules, is also due to inefficient removal of the cohesin from the arm regions and hence the resolution of chiasmata as we observed that removal of chiasmata could rescue the defect of prolonged anaphase I (Fig. S4A).

In *S. cerevisiae*, Rec8 removal from the chromatin is achieved by the protease separase that is released due to degradation of securin, Pds1. However, in *cin8*∆ *kip3*∆ cells, we observed uncoupling of Pds1 degradation from Rec8 removal (Fig. 5A-D). Thus it is reasonable to propose that in meiosis efficient Rec8 cleavage perhaps requires additional factor besides the release of separase.

### Microtubule based tension: a novel determinant to cleave Rec8 - but not Mcd1-cohesin?

It is important to address why Rec8 removal is compromised in absence of Cin8 and Kip3 together. Both Cin8 and Kip3 localize at the kinetochore where Kip3 is a part of the core kinetochore and is involved in kinetochore-microtubule attachment (Tytell and Sorger, 2006). On the other hand lack of both Cin8 and Kip3, but not individually, causes reduced transient separation of the sister kinetochores over the wild-type in the pre-anaphase mitotic cells (Tytell and Sorger, 2006) and we noticed that metaphase to anaphase transition is delayed (Fig. S2C). These results suggest that Cin8 and Kip3 together are involved in force generation on the chromosomes towards the opposite spindle poles which is consistent with the fact that these motors have microtubule cross-linking (Gordon and Roof, 1999; Hildebrandt et al., 2006; Su et al., 2013a) and depolymerase activities (Gardner et al., 2008; Su et al., 2013b; Su et al., 2011b). Therefore, in *cin8*∆ *kip3*∆ meiotic cells the homologs are not under tension in meiosis I and so are the sisters in meiosis II. We propose a model (Fig. 10) where the efficient cleavage of Rec8 both in meiosis I and II requires that the homologs and the sisters, respectively must be under tension. In support of this, in a phosphodeficient Cin8 mutant that remains bound to the spindle for longer and can generate force, the Rec8 removal is supposedly better and hence we observed less chromosome breakage and improved spore viability which was found opposite in case of a phosphomimic mutant that fails to bind to the spindle and generate force (Fig. 8B-C). To reconfirm our tension model of Rec8 cleavage, we created tensionless condition by removing either chiasmata (*SPO11*) or microtubules in a Mad2 depleted strains and observed defective Rec8 removal in both the conditions (Fig. 9A and D). Importantly, we failed to observe any perturbation in Mcd1 removal in the *cin8*∆ *kip3*∆ cells (Fig. 5E) and believe that is why the cells perform better in mitosis (Fig. S2D-E). To address how tension might drive Rec8 but not Mcd1 cleavage, we reason that the tension by some means may promote phosphorylation of Rec8 and earlier studies have shown that phosphorylation of Rec8, but not Mcd1, is indispensable for cohesin cleavage (Alexandru et al., 2001; Attner et al., 2013; Lee and Amon, 2003). The tension can influence maintenance of certain proteins on the chromatin responsible for Rec8 phosphorylation or a direct effect of tension on the cohesin may expose the Rec8 sites for phosphorylation. Alternatively, the tension may facilitate phosphorylated Rec8 amenable to cleavage by separase. It is possible that in *cin8*∆ *kip3*∆ cells, the spindle assembly checkpoint becomes activated as there is loss of tension and faulty kinetochore-microtubule attachment which can keep APC inactivated and resist cohesin cleavage. However, we believe this is unlikely as we observed Pds1 degradation occurred in the double mutant at a normal pace of the cell cycle (Fig. 5B and D). To explain why a tension based mechanism has evolved to sensitize cohesin removal in meiosis, it can be argued that in this cell cycle, unlike mitosis, chiasmata are formed and removal of arm cohesin is required for their resolution. During resolution, the ‘terminalization’ of the cross-over point that occurs due to pulling of the homologs might subject arms cohesin under tension that perhaps signals their removal. However, how prolonged retention of cohesin with reduced tension acting on the chromosomes (due to absence of Cin8 and Kip3) can eventually lead to chromosome breakage is not clear from our study. We observed that in the *cin8*∆ *kip3*∆ cells spindle disassembly is delayed (Fig. 1C) and Kip1 activity is protracted (Fig.7). Additionally, loss of these proteins can potentially cause abnormally extended kinetochore-microtubules since they also possess the microtubule depolymerase activity (Gardner et al., 2008; Su et al., 2013a; Su et al., 2011a). It is plausible that when these conditions together prevail over an extended period of time spanning two rounds of spindle assembly/disassembly and chromosome movement in meiosis, an imbalance of force is generated on the chromosomes causing them to break. In summary, we report here for the first time about the requirement of tension in the efficient removal of cohesin in meiosis and the importance of kinesin-5 and kinesin-8 motors in promoting this event and thus maintaining chromosome integrity. Given meiosis and the functions of kinesins are conserved across eukaryotes, it would be tempting to investigate if an attenuation of motor functions could be one of the reasons to generate aneuploid gametes that occurs at an alarming rate during human gametogenesis.

## Experimental procedures

### Yeast strains and Media

All the strains used in this study were of SK1 background. The list of the strains and plasmids with the genotype is mentioned in Table S1.The plasmids utilized for the C-terminal protein tagging and deletion of a gene are from Euroscarf and were PCR based (Wach et al., 1997). Transformation of the cells with the PCR cassettes was performed as mentioned earlier (Gietz and Schiestl, 2007). In case of selecting the cells on a dropout media along with the antibiotic G418, the media was used as mentioned earlier (Cheng et al., 2000) where instead of ammonium sulfate, monosodium glutamate was used to restore the sensitivity of G418. For metaphase I and prophase I arrest, *P_CLB2_* and *P_GAL1_* constructs were used to shuffle the endogenous promoters of *CDC20* and *NDT80*, respectively as mentioned earlier (Benjamin et al., 2003; Lee and Amon, 2003). For chromosome segregation assay, chromosome V and chromosome III were marked with GFP by integrating repeats of *tet* operators at 1.4 kb and *lac* operators at 22 kb away from the centromeres, respectively in a cell expressing TetR-GFP and LacI-GFP, respectively (Straight et al., 1996; Tanaka et al., 2000).

### Fluorescence microscopy

For live cell imaging, 1ml of 1 O.D_600_ culture was fixed with formaldehyde (final concentration 5%) for 5-10 min. The pellet was washed two times with 0.1M phosphate buffer (pH 7.5). For DAPI staining, after fixation of the sample with formaldehyde, the pellet was washed once with 50% ethanol and then resuspended in DAPI with the final concentration of 1µg/ml. The image was acquired with z-stacking (spacing 0.5 µm) using AxioObserver Z1 Zeiss inverted microscope (63X 1.4 NA Objective). Processing and merging of images were done using AxioVs40 V 4.8.2.0 software. Exposure time was set according to the fluorescence signal and was kept constant among the samples used for comparison (mainly it was 1.5s for EGFP, CFP, and mcherry fluorophores excitation). In order to avoid the bleed-through of intense Spc42-CFP signal though GFP channel, Zeiss confocal laser scanning microscope (LSM 780) equipped with 32 array GaAsP detector was used. Images were acquired using Zeiss Zen 2012 software.

### Image analysis

Images were generated by merging the planes projecting maximum intensity and were further analyzed. The quantification of fluorescent intensity of the images acquired using Zeiss Axio observerZ1 was performed using ImageJ software. A region of interest covering the fluorescence signal was defined and the integrated intensity of that region was estimated, following background reduction, by averaging the integrated intensity of three random non-fluorescent areas multiplied by the area of the fluorescent signal region. Estimation of number of Rad52 or Rec8 foci per chromosome spread was performed using an automatic spot detection algorithm (Imaris3D reconstitution software), keeping the threshold limit constant for all the images.

### Growth conditions and meiotic induction

Before meiotic induction, the cells were patched on YPG (yeast extract 1% peptone 2%, glycerol 2%) to restrain the growth of the petite colonies and then were transferred to presporulation medium for overnight. This was followed by meiotic induction in sporulation medium (0.02% raffinose, 1% potassium acetate) as mentioned elsewhere (Cha et al., 2000; Mehta et al., 2014).

In order to prevent the loss of centromeric plasmid containing mutated ORF of *CIN8* in the presporulation medium (PSP2) used for the meiosis synchronization, instead of yeast extract potassium acetate (YPA) medium, selective medium (SC–Uradropout) supplemented with 0.1% yeast extract was used (Roth and Halvorson, 1969; Simchen et al., 1972).

For mitotic synchronization, the cells were arrested at G1 using α-factor at a concentration of 5 µg/ml in 0.3 O.D_600_ of cells (Cui et al., 2009; Prajapati et al., 2017). After 3 h when >90% cells exhibited the shmoo formation, they were washed and released into fresh YPD medium.

For enhancing the expression of LacI-GFP that is under the control of the *HIS3* promoter, 3-aminotriazole was added at the final concentration of 20 mM in the sporulation medium.

Unlike mitosis, microtubule disruption before or during meiotic S phase causes cells to arrest at G1 or G2 phase, respectively (Hochwagen et al., 2005). Therefore, for microtubule depolymerization in meiosis, the cells were treated with benomyl at a concentration of 60 µg/ml after 5.5 h of meiotic induction when most of the cells passed through S phase. Stages of the cell cycle and microtubule morphology before and after the drug treatment were determined by tubulin immunofluorescence. In order to avoid the spindle checkpoint mediated arrest in the absence of microtubule, Mad2 was depleted in meiosis using *CLB2* promoter (Jin et al., 2009).

### Comet assay

Comet assay was performed as mentioned earlier (Oliveira and Johansson, 2012). After the meiotic induction in the SPM for 8h, 1 ml of sporulating culture with a concentration of 10^7^ cells per ml was harvested. Cells were then resuspended in the buffer (1 M sorbitol, 25 mM KH_2_PO_4_, 50 mM β-mercaptoethanol) containing Zymolase −20T (20mg/ml - MP biomedicals). pH of the buffer was adjusted to 6.5 using NaOH. The cells were then incubated for half an hour for making spheroplasts. Spheroplasted cells were then mixed with 1.5% low-melting-point (LMP) agarose and spreaded immediately on the glass slide precoated with 0.5% normal melting point (NMP) agarose. Slides were placed on ice for agarose to solidify for which the embedded cells form cavities in the gel. Subsequently, the slides were submerged into lysing buffer (30 mM NaOH, 1 M NaCl, 0.05% sodium dodecyl sulfate, 50 mM EDTA, 10 mM Tris-HCl, pH 10) for 20 min at 4°C. Following lysis of the spheroplasts, the cavities formed by the spheroplasted cells contained only high molecular weight DNA while the other biomolecules diffused out. The slides were then placed in the electrophoresis buffer (30 mM NaOH, 10 mM EDTA, 10 mM Tris-HCl, pH 10) at 4°C for 20 min for the unwinding of the DNA which was followed by electrophoresis for 20 min at 0.7V/cm. On application of the electric current of 300 mA at 24V, the fragmented DNA, named as the ‘tail’, moved towards the anode (+) while the compact mass of DNA remained in the cavity giving a ‘comet’-like appearance on the gel. Following this, the slides were incubated into the neutralization buffer (10 mM Tris-HCl, pH 7.4) at room temperature for 10 min. The slides were incubated in 76% and 96% ethanol for 10 min each at room temperature. The slides were then incubated with solution containing ethidium bromide (10µg/ml) for 5 min and were observed using epifluorescence microscope (excitation filter 546 nm; emission filter 575 nm). The wild-type cells treated with 10 mM concentration of H_2_O_2_ were used as positive control.

### Immunostaining

Immunostaining was performed as described earlier (Mehta et al., 2014). The cells from the meiotic culture were harvested and fixed with 5% formaldehyde. The spheroplasts were made using zymolyase, and were placed on a polylysine-coated slide. The spheroplasts were permeabilized by Triton X-100 or methanol/acetone and were then incubated with primary followed by secondary antibody. DAPI (4’, 6-diamidino-2-phenylindole) at a concentration of 1 μg/ml in 0.1 M phosphate buffer was used to stain the DNA. Primary antibodies including rat anti-tubulin (MCA78G; Serotec) and mouse anti-Myc (11667149001; Roche) were used at a dilution of 1:5000 and 1:200, respectively. The secondary antibodies used from Jackson were TRITC goat anti-rat (115-485-166), Alexa-fluor 488 goat anti-rat (112-545-167), and TRITC goat anti-mouse (115-025-166) at a dilution of 1:200.

### Chromosome spread

Protocol for chromosome spread formation was followed as mentioned elsewhere (Mehta et al., 2014; Prajapati et al., 2018). 2 ml of meiotic culture was spheroplasted using 20T zymolase (10mg/ml) for 1 h with 1.42 M β-ME. The reaction was stopped by addition of 200 µl stop solution (0.1 M MES, 1 mM EDTA, 0.5 mM MgCl_2_, 1 M sorbitol, pH 6.4). Spheroplasted cells were fixed on acid washed slides with paraformaldehyde solution (4% paraformaldehyde, 3.4% sucrose with 2 drops of NaOH to dissolve paraformaldehyde) followed by addition of 1% lipsol to burst the cells. The slides were kept to dry overnight at room temperature after homogenously smearing the spheroplasts on the slide. Next day the slides were washed with 2 ml of 0.4% photoflow-200 (Kodak) followed by washing in phosphate buffer saline for 10 minutes. Before addition of primary antibodies, 100 µl of blocking solution (5% skim milk) was added to the slide for 30 minutes. Primary antibodies were diluted in PBS supplemented with 0.1% BSA (bovine serum albumin). Primary antibodies used were rabbit anti-Zip1 (SC 33733; Santa Cruz Biotechnology, 1:100), mouse anti-HA (MMS-101P; Covance, 1:200), mouse anti-GFP (11814460001; Roche, 1:200). Slides are coated with 100 µl of primary antibody for 1 hour followed by washing with PBS three times with 5 min incubation each time. Similar treatment with secondary antibody was performed. Jackson secondary antibodies - TRITC goat anti-rat and Alexa-fluor 488 goat anti-mouse (115-485-166) were used at the dilution of 1:200. Chromatin was stained using DAPI.

### Immunoblotting and its quantification

Whole cell proteins were extracted by NaOH treatment as described earlier (Kushnirov, 2000) with some modifications. Cells from 10 ml of 1 O.D_600_ culture were pelleted down and treated with 0.1N NaOH for 30 min. After alkaline treatment, pelleted cells were resuspended in electrophoresis sample buffer (ESB; 2% SDS, 10% glycerol, 80 mM Tris pH 6.8, 2% bromophenol blue, 100 mM dithiothreitol; Dunn, 1986) and boiled for 5 min at 100°C. The supernatant obtained after the centrifugation was used for the immunoblotting. Primary antibody rabbit anti-Myc (ab9106; Abcam) was used at the dilution of 1:5000 in 1:20 TBST:5% skim milk. Jackson HRP conjugated secondary antibodies used for the detection were goat anti-mouse (115-035-166; 1:5000), goat anti-rabbit (111-035-003; 1:10000), and goat anti-rat (112-035-167; 1:10000). Blots were developed using ECL reagents (170-5060; Bio-Rad laboratories). The intensities of the bands at different time points were quantified using ImageJ software. The ratio of the protein bands to the loading control band was used for the comparison between the wild-type and mutant strains.

## Acknowledgement

We are grateful to Leah Gheber for providing the *cin8* phosphomutant plasmids. We thank the central instrumental facility of IIT Bombay for the Laser scanning confocal microscope. SKG lab is supported by DBT (BT/PR13909/BRB/10/1432/2015 and BT/PR20932/BRB/10/1539/2016) and CSIR (38(1457)/18/EMR-II) grants. PM is funded by UGC fellowship (17-06/2012(i) EU-V).

## Reference

Agarwal, M., G. Mehta, and S.K. Ghosh. 2015. Role of Ctf3 and COMA subcomplexes in meiosis: Implication in maintaining Cse4 at the centromere and numeric spindle poles. Biochim Biophys Acta. 1853:671–684.

Alexandru, G., F. Uhlmann, K. Mechtler, M.A. Poupart, and K. Nasmyth. 2001. Phosphorylation of the cohesin subunit Scc1 by Polo/Cdc5 kinase regulates sister chromatid separation in yeast. Cell. 105:459–472.

Attner, M.A., M.P. Miller, L.-s. Ee, S. K. Elkin, and A. Amon. 2013. Polo kinase Cdc5 is a central regulator of meiosis I. Proceedings of the National Academy of Sciences of the United States of America. 110:14278–14283.

Avunie-Masala, R., N. Movshovich, Y. Nissenkorn, A. Gerson-Gurwitz, V. Fridman, M. Kõivomägi, M. Loog, M.A. Hoyt, A. Zaritsky, and L. Gheber. 2011. Phospho-regulation of kinesin-5 during anaphase spindle elongation. Journal of cell science. 124:873–878.

Barlow, J.H., and R. Rothstein. 2009. Rad52 recruitment is DNA replication independent and regulated by Cdc28 and the Mec1 kinase. The EMBO Journal. 28:1121–1130.

Barton, N.R., and L.S. Goldstein. 1996. Going mobile: microtubule motors and chromosome segregation. Proceedings of the National Academy of Sciences of the United States of America. 93:1735–1742.

Bascom-Slack, C.A., and D.S. Dawson. 1997. The Yeast Motor Protein, Kar3p, Is Essential for Meiosis I. The Journal of Cell Biology. 139:459–467.

Benjamin, K.R., C. Zhang, K.M. Shokat, and I. Herskowitz. 2003. Control of landmark events in meiosis by the CDK Cdc28 and the meiosis-specific kinase Ime2. Genes Dev. 17:1524–1539.

Buonomo, S.B., R.K. Clyne, J. Fuchs, J. Loidl, F. Uhlmann, and K. Nasmyth. 2000. Disjunction of homologous chromosomes in meiosis I depends on proteolytic cleavage of the meiotic cohesin Rec8 by separin. Cell. 103:387–398.

Buonomo, S.B., K.P. Rabitsch, J. Fuchs, S. Gruber, M. Sullivan, F. Uhlmann, M. Petronczki, A. Toth, and K. Nasmyth. 2003. Division of the nucleolus and its release of CDC14 during anaphase of meiosis I depends on separase, SPO12, and SLK19. Developmental cell. 4:727–739.

Cha, R.S., B.M. Weiner, S. Keeney, J. Dekker, and N. Kleckner. 2000. Progression of meiotic DNA replication is modulated by interchromosomal interaction proteins, negatively by Spo11p and positively by Rec8p. Genes Dev. 14:493–503.

Challa, K., V.G. Fajish, M. Shinohara, F. Klein, S.M. Gasser, and A. Shinohara. 2019. Meiosis-specific prophase-like pathway controls cleavage-independent release of cohesin by Wapl phosphorylation. 15:e1007851.

Chee, M.K., and S.B. Haase. 2010. B-cyclin/CDKs regulate mitotic spindle assembly by phosphorylating kinesins-5 in budding yeast. PLoS genetics. 6:e1000935.

Chen, J.-S., M.R. Broadus, J.R. McLean, A. Feoktistova, L. Ren, and K.L. Gould. 2013. Comprehensive Proteomics Analysis Reveals New Substrates and Regulators of the Fission Yeast Clp1/Cdc14 Phosphatase. Molecular & Cellular Proteomics. 12:1074–1086.

Cheng, T.H., C.R. Chang, P. Joy, S. Yablok, and M.R. Gartenberg. 2000. Controlling gene expression in yeast by inducible site-specific recombination. Nucleic acids research. 28:E108.

Conrad, M.N., C.Y. Lee, G. Chao, M. Shinohara, H. Kosaka, A. Shinohara, J.A. Conchello, and M.E. Dresser. 2008. Rapid telomere movement in meiotic prophase is promoted by NDJ1, MPS3, and CSM4 and is modulated by recombination. Cell. 133:1175–1187.

Cottingham, F.R., L. Gheber, D.L. Miller, and M.A. Hoyt. 1999. Novel Roles for Saccharomyces cerevisiae Mitotic Spindle Motors. The Journal of cell biology. 147:335–350.

D’Amours, D., and A. Amon. 2004. At the interface between signaling and executing anaphase--Cdc14 and the FEAR network. Genes Dev. 18:2581–2595.

Dagenbach, E.M., and S.A. Endow. 2004. A new kinesin tree. Journal of cell science. 117:3–7.

DeZwaan, T.M., E. Ellingson, D. Pellman, and D.M. Roof. 1997. Kinesin-related KIP3 of Saccharomyces cerevisiae is required for a distinct step in nuclear migration. The Journal of cell biology. 138:1023–1040.

Dunn, S.D. 1986. Effects of the modification of transfer buffer composition and the renaturation of proteins in gels on the recognition of proteins on Western blots by monoclonal antibodies. Analytical biochemistry. 157:144–153.

Fridman, V., A. Gerson-Gurwitz, O. Shapira, N. Movshovich, S. Lakamper, C.F. Schmidt, and L. Gheber. 2013. Kinesin-5 Kip1 is a bi-directional motor that stabilizes microtubules and tracks their plus-ends in vivo. Journal of cell science. 126:4147–4159.

Friel, C.T., and J. Howard. 2011. The kinesin-13 MCAK has an unconventional ATPase cycle adapted for microtubule depolymerization. The EMBO Journal. 30:3928–3939.

Gardner, M.K., D.C. Bouck, L.V. Paliulis, J.B. Meehl, E.T. O’Toole, J. Haase, A. Soubry, A.P. Joglekar, M. Winey, E.D. Salmon, K. Bloom, and D.J. Odde. 2008. Chromosome Congression by Kinesin-5 Motor-Mediated Disassembly of Longer Kinetochore Microtubules. Cell. 135:894–906.

Gasior, S.L., A.K. Wong, Y. Kora, A. Shinohara, and D.K. Bishop. 1998. Rad52 associates with RPA and functions with Rad55 and Rad57 to assemble meiotic recombination complexes. Genes & Development. 12:2208–2221.

Geiser, J.R., E.J. Schott, T.J. Kingsbury, N.B. Cole, L.J. Totis, G. Bhattacharyya, L. He, and M.A. Hoyt. 1997. Saccharomyces cerevisiae genes required in the absence of the CIN8-encoded spindle motor act in functionally diverse mitotic pathways. Mol Biol Cell. 8:1035–1050.

Gerson-Gurwitz, A., C. Thiede, N. Movshovich, V. Fridman, M. Podolskaya, T. Danieli, S. Lakämper, D.R. Klopfenstein, C.F. Schmidt, and L. Gheber. 2011. Directionality of individual kinesin-5 Cin8 motors is modulated by loop 8, ionic strength and microtubule geometry. The EMBO Journal. 30:4942–4954.

Gietz, R.D., and R.H. Schiestl. 2007. High-efficiency yeast transformation using the LiAc/SS carrier DNA/PEG method. Nature protocols. 2:31–34.

Gladstone, M.N., D. Obeso, H. Chuong, and D.S. Dawson. 2009. The synaptonemal complex protein Zip1 promotes bi-orientation of centromeres at meiosis I. PLoS Genet. 5:e1000771.

Goldstein, A., D. Goldman, E. Valk, M. Loog, L.J. Holt, and L. Gheber. 2018. Synthetic-evolution reveals that phosphoregulation of the mitotic kinesin-5 Cin8 is constrained. bioRxiv:312637.

Goldstein, A., N. Siegler, D. Goldman, H. Judah, E. Valk, M. Kõivomägi, M. Loog, and L. Gheber. 2017. Three Cdk1 sites in the kinesin-5 Cin8 catalytic domain coordinate motor localization and activity during anaphase. Cellular and Molecular Life Sciences. 74:3395–3412.

Gordon, D.M., and D.M. Roof. 1999. The kinesin-related protein Kip1p of Saccharomyces cerevisiae is bipolar. The Journal of biological chemistry. 274:28779–28786.

Gordon, D.M., and D.M. Roof. 2001. Degradation of the kinesin Kip1p at anaphase onset is mediated by the anaphase-promoting complex and Cdc20p. Proceedings of the National Academy of Sciences. 98:12515–12520.

Gupta Jr, M.L., P. Carvalho, D.M. Roof, and D. Pellman. 2006. Plus end-specific depolymerase activity of Kip3, a kinesin-8 protein, explains its role in positioning the yeast mitotic spindle. Nature cell biology. 8:913.

Gupta, M.L., Jr., P. Carvalho, D.M. Roof, and D. Pellman. 2006. Plus end-specific depolymerase activity of Kip3, a kinesin-8 protein, explains its role in positioning the yeast mitotic spindle. Nature cell biology. 8:913–923.

Hildebrandt, E.R., L. Gheber, T. Kingsbury, and M.A. Hoyt. 2006. Homotetrameric form of Cin8p, a Saccharomyces cerevisiae kinesin-5 motor, is essential for its in vivo function. The Journal of biological chemistry. 281:26004–26013.

Hildebrandt, E.R., and M.A. Hoyt. 2000. Mitotic motors in Saccharomyces cerevisiae. Biochim Biophys Acta. 1496:99–116.

Hildebrandt, E.R., and M.A. Hoyt. 2001. Cell cycle-dependent degradation of the Saccharomyces cerevisiae spindle motor Cin8p requires APC(Cdh1) and a bipartite destruction sequence. Mol Biol Cell. 12:3402–3416.

Hochwagen, A., G. Wrobel, M. Cartron, P. Demougin, C. Niederhauser-Wiederkehr, M.G. Boselli, M. Primig, and A. Amon. 2005. Novel Response to Microtubule Perturbation in Meiosis. Molecular and Cellular Biology. 25:4767–4781.

Hoyt, M.A., L. He, K.K. Loo, and W.S. Saunders. 1992a. Two Saccharomyces cerevisiae kinesin-related gene products required for mitotic spindle assembly. The Journal of cell biology. 118:109–120.

Hoyt, M.A., L. He, K.K. Loo, and W.S. Saunders. 1992b. Two Saccharomyces cerevisiae kinesin-related gene products required for mitotic spindle assembly. The Journal of Cell Biology. 118:109–120.

Hoyt, M.A., L. He, L. Totis, and W.S. Saunders. 1993. Loss of Function of Saccharomyces Cerevisiae Kinesin-Related Cin8 and Kip1 Is Suppressed by Kar3 Motor Domain Mutations. Genetics. 135:35–44.

Hunter, A.W., M. Caplow, D.L. Coy, W.O. Hancock, S. Diez, L. Wordeman, and J. Howard. 2003. The kinesin-related protein MCAK is a microtubule depolymerase that forms an ATP-hydrolyzing complex at microtubule ends. Molecular cell. 11:445–457.

Jin, H., V. Guacci, and H.-G. Yu. 2009. Pds5 is required for homologue pairing and inhibits synapsis of sister chromatids during yeast meiosis. The Journal of cell biology. 186:713–725.

Kamieniecki, R.J., R.M.Q. Shanks, and D.S. Dawson. 2000. Slk19p is necessary to prevent separation of sister chromatids in meiosis I. Current Biology. 10:1182–1190.

Kapitein, L.C., E.J.G. Peterman, B.H. Kwok, J.H. Kim, T.M. Kapoor, and C.F. Schmidt. 2005. The bipolar mitotic kinesin Eg5 moves on both microtubules that it crosslinks. Nature. 435:114.

Kashlna, A.S., R.J. Baskin, D.G. Cole, K.P. Wedaman, W.M. Saxton, and J.M. Scholey. 1996. A bipolar kinesin. Nature. 379:270.

Katis, V.L., M. Galova, K.P. Rabitsch, J. Gregan, and K. Nasmyth. 2004. Maintenance of Cohesin at Centromeres after Meiosis I in Budding Yeast Requires a Kinetochore-Associated Protein Related to MEI-S332. Current Biology. 14:560–572.

Khmelinskii, A., and E. Schiebel. 2008. Assembling the spindle midzone in the right place at the right time. Cell cycle (Georgetown, Tex.). 7:283–286.

Klapholz, S., C.S. Waddell, and R.E. Esposito. 1985. The role of the SPO11 gene in meiotic recombination in yeast. Genetics. 110:187–216.

Klein, F., P. Mahr, M. Galova, S.B. Buonomo, C. Michaelis, K. Nairz, and K. Nasmyth. 1999. A central role for cohesins in sister chromatid cohesion, formation of axial elements, and recombination during yeast meiosis. Cell. 98:91–103.

Koszul, R., K.P. Kim, M. Prentiss, N. Kleckner, and S. Kameoka. 2008. Meiotic chromosomes move by linkage to dynamic actin cables with transduction of force through the nuclear envelope. Cell. 133:1188–1201.

Kushnirov, V.V. 2000. Rapid and reliable protein extraction from yeast. Yeast (Chichester, England). 16:857–860.

Lee, B.H., and A. Amon. 2003. Role of Polo-like kinase CDC5 in programming meiosis I chromosome segregation. Science (New York, N.Y.). 300:482–486.

Malone, R.E., and R.E. Esposito. 1980. The RAD52 gene is required for homothallic interconversion of mating types and spontaneous mitotic recombination in yeast. Proceedings of the National Academy of Sciences of the United States of America. 77:503–507.

Marston, A.L., B.H. Lee, and A. Amon. 2003. The Cdc14 phosphatase and the FEAR network control meiotic spindle disassembly and chromosome segregation. Developmental cell. 4:711–726.

Mayer, M.L., I. Pot, M. Chang, H. Xu, V. Aneliunas, T. Kwok, R. Newitt, R. Aebersold, C. Boone, G.W. Brown, and P. Hieter. 2004. Identification of Protein Complexes Required for Efficient Sister Chromatid Cohesion. Molecular Biology of the Cell. 15:1736–1745.

Mayr, M.I., M. Storch, J. Howard, and T.U. Mayer. 2011. A non-motor microtubule binding site is essential for the high processivity and mitotic function of kinesin-8 Kif18A. PloS one. 6:e27471.

Mehta, G.D., M. Agarwal, and S.K. Ghosh. 2014. Functional characterization of kinetochore protein, Ctf19 in meiosis I: an implication of differential impact of Ctf19 on the assembly of mitotic and meiotic kinetochores in Saccharomyces cerevisiae. Mol Microbiol. 91:1179–1199.

Miloshev, G., I. Mihaylov, and B. Anachkova. 2002. Application of the single cell gel electrophoresis on yeast cells. Mutation research. 513:69–74.

Nerusheva, O.O., S. Galander, J. Fernius, D. Kelly, and A.L. Marston. 2014. Tension-dependent removal of pericentromeric shugoshin is an indicator of sister chromosome biorientation. Genes & development. 28:1291–1309.

Oliveira, R., and B. Johansson. 2012. Quantitative DNA damage and repair measurement with the yeast comet assay. Methods in molecular biology (Clifton, N.J.). 920:101–109.

Orr-Weaver, T.L., J.W. Szostak, and R.J. Rothstein. 1981. Yeast transformation: a model system for the study of recombination. Proc Natl Acad Sci U S A. 78:6354–6358.

Ostling, O., and K.J. Johanson. 1984. Microelectrophoretic study of radiation-induced DNA damages in individual mammalian cells. Biochemical and biophysical research communications. 123:291–298.

Penkner, A.M., A. Fridkin, J. Gloggnitzer, A. Baudrimont, T. Machacek, A. Woglar, E. Csaszar, P. Pasierbek, G. Ammerer, Y. Gruenbaum, and V. Jantsch. 2009. Meiotic Chromosome Homology Search Involves Modifications of the Nuclear Envelope Protein Matefin/SUN-1. Cell. 139:920–933.

Prajapati, H.K., M. Agarwal, P. Mittal, and S.K. Ghosh. 2018. Evidence of Zip1 Promoting Sister Kinetochore Mono-orientation During Meiosis in Budding Yeast. G3: Genes|Genomes|Genetics. 8:3691–3701.

Queralt, E., C. Lehane, B. Novak, and F. Uhlmann. 2006. Downregulation of PP2A(Cdc55) phosphatase by separase initiates mitotic exit in budding yeast. Cell. 125:719–732.

Resnick, M.A., and P. Martin. 1976. The repair of double-strand breaks in the nuclear DNA of Saccharomyces cerevisiae and its genetic control. Molecular & general genetics: MGG. 143:119–129.

Riedel, C.G., V.L. Katis, Y. Katou, S. Mori, T. Itoh, W. Helmhart, M. Galova, M. Petronczki, J. Gregan, B. Cetin, I. Mudrak, E. Ogris, K. Mechtler, L. Pelletier, F. Buchholz, K. Shirahige, and K. Nasmyth. 2006. Protein phosphatase 2A protects centromeric sister chromatid cohesion during meiosis I. Nature. 441:53–61.

Rizk, R.S., K.A. DiScipio, K.G. Proudfoot, and M.L. Gupta. 2014. The kinesin-8 Kip3 scales anaphase spindle length by suppression of midzone microtubule polymerization. The Journal of cell biology. 204:965–975.

Roccuzzo, M., C. Visintin, F. Tili, and R. Visintin. 2015. FEAR-mediated activation of Cdc14 is the limiting step for spindle elongation and anaphase progression. Nature cell biology. 17:251–261.

Rock, J.M., and A. Amon. 2009. The FEAR network. Current biology: CB. 19: 10.1016/j.cub.2009.1010.1002.

Roof, D.M., P.B. Meluh, and M.D. Rose. 1992. Kinesin-related proteins required for assembly of the mitotic spindle. The Journal of cell biology. 118:95.

Roostalu, J., C. Hentrich, P. Bieling, I.A. Telley, E. Schiebel, and T. Surrey. 2011. Directional Switching of the Kinesin Cin8 Through Motor Coupling. Science (New York, N.Y.).

Roth, R., and H.O. Halvorson. 1969. Sporulation of Yeast Harvested During Logarithmic Growth. Journal of Bacteriology. 98:831–832.

Sato, A., B. Isaac, C.M. Phillips, R. Rillo, P.M. Carlton, D.J. Wynne, R.A. Kasad, and A.F. Dernburg. 2009. Cytoskeletal Forces Span the Nuclear Envelope to Coordinate Meiotic Chromosome Pairing and Synapsis. Cell. 139:907–919.

Saunders, W., D. Hornack, V. Lengyel, and C. Deng. 1997. The Saccharomyces cerevisiae Kinesin-related Motor Kar3p Acts at Preanaphase Spindle Poles to Limit the Number and Length of Cytoplasmic Microtubules. The Journal of Cell Biology. 137:417–431.

Saunders, W.S., and M.A. Hoyt. 1992. Kinesin-related proteins required for structural integrity of the mitotic spindle. Cell. 70:451–458.

Saunders, W.S., D. Koshland, D. Eshel, I.R. Gibbons, and M.A. Hoyt. 1995. Saccharomyces cerevisiae kinesin- and dynein-related proteins required for anaphase chromosome segregation. The Journal of cell biology. 128:617.

Shanks, R.M., R.J. Kamieniecki, and D.S. Dawson. 2001. The Kar3-interacting protein Cik1p plays a critical role in passage through meiosis I in Saccharomyces cerevisiae. Genetics. 159:939–951.

Shapira, O., and L. Gheber. 2016. Motile properties of the bi-directional kinesin-5 Cin8 are affected by phosphorylation in its motor domain. Scientific reports. 6:25597–25597.

Shapira, O., A. Goldstein, J. Al-Bassam, and L. Gheber. 2017. A potential physiological role for bi-directional motility and motor clustering of mitotic kinesin-5 Cin8 in yeast mitosis. 130:725–734.

Shonn, M.A., R. McCarroll, and A.W. Murray. 2002. Spo13 protects meiotic cohesin at centromeres in meiosis I. Genes & development. 16:1659–1671.

Simchen, G., R. Pinon, and Y. Salts. 1972. Sporulation in Saccharomyces cerevisiae: premeiotic DNA synthesis, readiness and commitment. Experimental cell research. 75:207–218.

Singh, S.K., H. Pandey, J. Al-Bassam, and L. Gheber. 2018. Bidirectional motility of kinesin-5 motor proteins: structural determinants, cumulative functions and physiological roles. Cellular and Molecular Life Sciences. 75:1757–1771.

Stegmeier, F., R. Visintin, and A. Amon. 2002. Separase, polo kinase, the kinetochore protein Slk19, and Spo12 function in a network that controls Cdc14 localization during early anaphase. Cell. 108:207–220.

Straight, A.F., A.S. Belmont, C.C. Robinett, and A.W. Murray. 1996. GFP tagging of budding yeast chromosomes reveals that protein-protein interactions can mediate sister chromatid cohesion. Current biology: CB. 6:1599–1608.

Su, X., H. Arellano-Santoyo, D. Portran, J. Gaillard, M. Vantard, M. Thery, and D. Pellman. 2013a. Microtubule-sliding activity of a kinesin-8 promotes spindle assembly and spindle-length control. Nature cell biology. 15:948–957.

Su, X., H. Arellano-Santoyo, D. Portran, J. Gaillard, M. Vantard, M. Thery, and D. Pellman. 2013b. Microtubule-sliding activity of a kinesin-8 promotes spindle assembly and spindle-length control. Nature cell biology. 15:948.

Su, X., W. Qiu, M.L. Gupta, Jr., J.B. Pereira-Leal, S.L. Reck-Peterson, and D. Pellman. 2011a. Mechanisms underlying the dual-mode regulation of microtubule dynamics by Kip3/kinesin-8. Molecular cell. 43:751–763.

Su, X., W. Qiu, Mohan L. Gupta, José B. Pereira-Leal, Samara L. Reck-Peterson, and D. Pellman. 2011b. Mechanisms Underlying the Dual-Mode Regulation of Microtubule Dynamics by Kip3/Kinesin-8. Molecular cell. 43:751–763.

Tanaka, K., E. Kitamura, Y. Kitamura, and T.U. Tanaka. 2007. Molecular mechanisms of microtubule-dependent kinetochore transport toward spindle poles. The Journal of cell biology. 178:269–281.

Tanaka, T., J. Fuchs, J. Loidl, and K. Nasmyth. 2000. Cohesin ensures bipolar attachment of microtubules to sister centromeres and resists their precocious separation. Nature cell biology. 2:492–499.

Tang, N.H., and T. Toda. 2015. Alp7/TACC recruits kinesin-8-PP1 to the Ndc80 kinetochore protein for timely mitotic progression and chromosome movement. Journal of cell science. 128:354–363.

Toth, A., K.P. Rabitsch, M. Galova, A. Schleiffer, S.B. Buonomo, and K. Nasmyth. 2000. Functional genomics identifies monopolin: a kinetochore protein required for segregation of homologs during meiosis i. Cell. 103:1155–1168.

Tytell, J.D., and P.K. Sorger. 2006. Analysis of kinesin motor function at budding yeast kinetochores. J Cell Biol. 172:861–874.

Ubersax, J.A., E.L. Woodbury, P.N. Quang, M. Paraz, J.D. Blethrow, K. Shah, K.M. Shokat, and D.O. Morgan. 2003. Targets of the cyclin-dependent kinase Cdk1. Nature. 425:859–864.

Wach, A., A. Brachat, C. Alberti-Segui, C. Rebischung, and P. Philippsen. 1997. Heterologous HIS3 marker and GFP reporter modules for PCR-targeting in Saccharomyces cerevisiae. Yeast (Chichester, England). 13:1065–1075.

Wanat, J.J., K.P. Kim, R. Koszul, S. Zanders, B. Weiner, N. Kleckner, and E. Alani. 2008. Csm4, in collaboration with Ndj1, mediates telomere-led chromosome dynamics and recombination during yeast meiosis. PLoS genetics. 4:e1000188.

Wells, J.L., D.W. Pryce, and R.J. McFarlane. 2006. Homologous chromosome pairing in Schizosaccharomyces pombe. Yeast (Chichester, England). 23:977–989.

Winey, M., and K. Bloom. 2012. Mitotic Spindle Form and Function. Genetics. 190:1197–1224.

Yamamoto, A., K. Kitamura, D. Hihara, Y. Hirose, S. Katsuyama, and Y. Hiraoka. 2008. Spindle checkpoint activation at meiosis I advances anaphase II onset via meiosis-specific APC/C regulation. The Journal of cell biology. 182:277–288.

Yellman, C.M., and G.S. Roeder. 2015. Cdc14 Early Anaphase Release, FEAR, Is Limited to the Nucleus and Dispensable for Efficient Mitotic Exit. PloS one. 10:e0128604.

Zeng, X., and W.S. Saunders. 2000. The *Saccharomyces cerevisiae* Centromere Protein Slk19p Is Required for Two Successive Divisions During Meiosis. Genetics. 155:577–587.

