## Supplemental file 1 for "Meiosis-specific functions of kinesin motors in cohesin removal and maintenance of chromosome integrity in budding yeast"

Table S1: **List of Sk1 background *S. cerevisiae* yeast strains used in this study**

| Yeast strain | Genotype |
|---|---|
| SGY40 | *MAT*a/α, *ho::LYS2, lys2, ura3, leu2::hisG, his3::hisG, trp1::hisG* |
| SGY155 | *MAT*a, *leu2::tetR-GFP::LEU2, CEN5::TetO-HIS3* |
| SGY263 | *MAT*a/α, *leu2::tetR-GFP::LEU2, CEN5::TetO-HIS3/leu2::tetRGFP::LEU2, CEN5::TetO-HIS3, ndt80::*$P_{GAL}$*-NDT80::TRP1/ ndt80::*$P_{GAL}$*-NDT80::TRP1* |
| SGY302 | *MAT*a, *leu2::tetR-GFP::LEU2, CEN5::TetO-HIS3, cin8Δ:: loxp-URA3-loxp* |
| SGY304 | *MAT*a, *leu2::tetR-GFP::LEU2, CEN5::TetO-HIS3, kip3Δ:: loxp-KanMx-loxp* |
| SGY309 | *MAT*a/α, *leu2::tetR-GFP::LEU2 CEN5::TetO-HIS3/leu2::tetRGFP, kip3Δ:: loxp-URA3-loxp/ kip3Δ:: loxp-URA3-loxp* |
| SGY314 | *MAT*a/α, *leu2::tetR-GFP::LEU2, CEN5::TetO-HIS3/leu2::tetRGFP::LEU2, CEN5::TetO-HIS3, kip3Δ:: loxp-URA3-loxp/ kip3Δ:: loxp-URA3-loxp* |
| SGY315 | *MAT*a/α, *leu2::tetR-GFP::LEU2, CEN5::TetO-HIS3/leu2::tetRGFP::LEU2, CEN5::TetO-HIS3, cin8Δ:: loxp-URA3-loxp/ cin8Δ:: loxp-URA3-loxp* |
| SGY317 | *MAT*a/α, *leu2::tetR-GFP::LEU2, CEN5::TetO-HIS3/leu2::tetRGFP::LEU2, CEN5::TetO-HIS3, kip1Δ:: loxp-URA3-loxp/ kip1Δ:: loxp-URA3-loxp* |
| SGY3248 | *MAT*a/α, *mad2::*$P_{CLB2}$*-MAD2::URA3/ mad2::*$P_{CLB2}$*-MAD2::URA3, REC8-EGFP:: HIS3/ REC8-EGFP:: HIS3, SPC42-CFP:: TRP1/ SPC42-CFP:: TRP1* |

Santanu K Ghosh

| | |
|---|---|
| SGY5001 | MATa/α, leu2::tetR-GFP::LEU2, CEN5::TetO-HIS3/leu2::tetRGFP::LEU2, CEN5::TetO-HIS3 |
| SGY5006 | MATa/α, leu2::tetR-GFP::LEU2 CEN5::TetO-HIS3/leu2::tetRGFP |
| SGY5009 | MATa/α, leu2::tetR-GFP::LEU2 CEN5::TetO-HIS3/leu2::tetRGFP::LEU2 CEN5::TetO-HIS3 kar3Δ:: loxp-URA3-loxp/ kar3Δ:: loxp-URA3-loxp |
| SGY5018 | MATa/α, leu2::tetR-GFP::LEU2 CEN5::TetO-HIS3/leu2::tetRGFP, cdc20::$P_{CLB2}$-CDC20::KanMx/ cdc20::$P_{CLB2}$-CDC20::KanMx, kip1Δ:: loxp-URA3-loxp/ kip1Δ:: loxp-URA3-loxp |
| SGY5034 | MATa/α, leu2::tetR-GFP::LEU2 CEN5::TetO-HIS3/leu2::tetRGFP, cdc20::$P_{CLB2}$-CDC20::KanMx/ cdc20::$P_{CLB2}$-CDC20::KanMx, kip3Δ:: loxp-URA3-loxp/ kip3Δ:: loxp-URA3-loxp |
| SGY5048 | MATa/α, leu2::tetR-GFP::LEU2, CEN5::TetO-HIS3/leu2::tetRGFP::LEU2 CEN5::TetO-HIS,3 spo11Δ:: loxp-URA3-loxp/ spo11Δ:: loxp-URA3-loxp |
| SGY5077 | MATa/α, leu2::tetR-GFP::LEU2 CEN5::TetO-HIS3/leu2::tetRGFP, kip1Δ:: loxp-URA3-loxp/ kip1Δ:: loxp-URA3-loxp |
| SGY5078 | MATa/α, leu2::tetR-GFP::LEU2 CEN5::TetO-HIS3/leu2::tetRGFP, cdc20::$P_{CLB2}$-CDC20::KanMx/ cdc20::$P_{CLB2}$-CDC20::KanMx, cin8Δ:: loxp-URA3-loxp/ cin8Δ:: loxp-URA3-loxp |
| SGY5089 | MATa/α, leu2::tetR-GFP::LEU2, CEN5::TetO-HIS3/leu2::tetRGFP::LEU2, CEN5::TetO-HIS3, cin8Δ:: loxp-URA3-loxp/ cin8Δ:: loxp-URA3-loxp, kip3Δ:: Loxp/ kip3Δ:: Loxp |
| SGY5090 | MATa/α, leu2::tetR-GFP::LEU2 CEN5::TetO-HIS3/leu2::tetRGFP, cin8Δ:: loxp-URA3-loxp/ cin8Δ:: loxp-URA3-loxp |
| SGY5104 | MATa/α, leu2::tetR-GFP::LEU2, CEN5::TetO-HIS3/leu2::tetRGFP::LEU2, CEN5::TetO-HIS3, kip1Δ:: loxp-URA3-loxp/ kip1Δ:: loxp-URA3-loxp, kip3Δ:: Loxp/ kip3Δ:: Loxp |
| SGY5124 | MATa, leu2::tetR-GFP::LEU2, CEN5::TetO-HIS3, cin8Δ:: loxp-URA3-loxp, kip3Δ:: Loxp |
| SGY5143 | MATa/α, leu2::tetR-GFP::LEU2 CEN5::TetO- |

Santanu K Ghosh

| | |
|---|---|
| | HIS3/leu2::tetRGFP, cdc20::$P_{CLB2}$-CDC20::KanMx/ cdc20::$P_{CLB2}$-CDC20::KanMx |
| SGY5154 | MATa/α, leu2::tetR-GFP::LEU2, CEN5::TetO-HIS3/leu2::tetRGFP::LEU2, cin8Δ:: loxp-URA3-loxp/ cin8Δ:: loxp-URA3-loxp, kip3Δ:: loxp-KanMx-loxp / kip3Δ:: loxp-KanMx-loxp |
| SGY5190 | MATa/α, leu2::tetR-GFP::LEU2, CEN5::TetO-HIS3/leu2::tetRGFP::LEU2, CEN5::TetO-HIS3, ndt80::$P_{GAL}$-NDT80::TRP1/ ndt80::$P_{GAL}$-NDT80::TRP1, kip1Δ:: loxp-URA3-loxp/ kip1Δ:: loxp-URA3-loxp |
| SGY5197 | MATa/α, leu2::tetR-GFP::LEU2, CEN5::TetO-HIS3/leu2::tetRGFP::LEU2,CEN5::TetO-HIS3, ndt80::$P_{GAL}$-NDT80::TRP1/ ndt80::$P_{GAL}$-NDT80::TRP1, cin8Δ:: loxp-KanMx-loxp/ cin8Δ:: loxp-KanMx-loxp |
| SGY5333 | MATa/α, NDC10-mCherry::KanMx/ NDC10-mCherry::KanMx, leu2::tetR-GFP::LEU2, CEN5::TetO-HIS3/leu2::tetRGFP::LEU2, cin8Δ:: Loxp/ cin8Δ:: Loxp, kip3Δ:: Loxp/ kip3Δ:: Loxp |
| SGY5329 | MATa/α, his3:GFP-LacI::HIS3, CEN5::LacO-LEU2/ his3:GFP-LacI::HIS3, CEN5::LacO-LEU2, cin8Δ:: loxp-URA3-loxp/ cin8Δ:: loxp-URA3-loxp, kip3Δ:: loxp-KanMx-loxp / kip3Δ:: loxp-KanMx-loxp |
| SGY5338 | MATa/α, SPC42-mCherry::KanMx/ SPC42-mCherry::KanMx, leu2::tetR-GFP::LEU2, CEN5::TetO-HIS3/leu2::tetRGFP::LEU2, cin8Δ:: Loxp/ cin8Δ:: Loxp, kip3Δ:: Loxp/ kip3Δ:: Loxp |
| SGY5347 | MATa/α, leu2::tetR-GFP::LEU2, CEN5::TetO-HIS3/leu2::tetRGFP::LEU2, CEN5::TetO-HIS3, ndt80::$P_{GAL}$-NDT80::TRP1/ ndt80::$P_{GAL}$-NDT80::TRP1, kip1Δ:: loxp-URA3-loxp/ kip1Δ:: loxp-URA3-loxp |
| SGY5357 | MATa, leu2::tetR-GFP::LEU2, CEN5::TetO-HIS3, cin8Δ:: loxp-URA3-loxp |
| SGY5385 | MATa/α, leu2::tetR-GFP::LEU2, CEN5::TetO-HIS3/leu2::tetRGFP::LEU2, CEN5::TetO-HIS3, cin8Δ:: loxp-URA3-loxp/ cin8Δ:: loxp-URA3-loxp, kip3Δ:: Loxp/ kip3Δ:: Loxp. ura3::CFP-TUB1::URA3/ ura3::CFP-TUB1::URA3 |
| SGY5407 | MATa/α, his3:GFP-LacI::HIS3, CEN5::LacO-LEU2/ his3:GFP-LacI::HIS3, CEN5::LacO-LEU2 |

Santanu K Ghosh

| | |
|---|---|
| SGY5414 | *MAT*a/α, *RAD52-EGFP::TRP1/ RAD52-EGFP::TRP1* |
| SGY5415 | *MAT*a/α, *RAD52-EGFP::TRP1/ RAD52-EGFP::TRP1, cin8Δ:: loxp-URA3-loxp/ cin8Δ:: loxp-URA3-loxp, kip3Δ:: loxp-URA3-loxp/ kip3Δ:: loxp-URA3-loxp* |
| SGY5422 | *MAT*a/α, *RAD52-EGFP::TRP1/ RAD52-EGFP::TRP1, cin8Δ:: Loxp/ cin8Δ:: Loxp* |
| SGY5444 | *MAT*a/α, *leu2::tetR-GFP::LEU2 CEN5::TetO-HIS3/leu2::tetRGFP::LEU2 CEN5::TetO-HIS,3 cin8Δ:: Loxp/ cin8Δ:: Loxp, kip3Δ:: Loxp/ kip3Δ:: Loxp, spo11Δ:: loxp-KanMx-loxp/ spo11Δ:: loxp-KanMx-loxp* |
| SGY5497 | *MAT*a/α, *REC8-EGFP::HIS3/ REC8-EGFP::HIS3* |
| SGY5500 | *MAT*a/α, *REC8-EGFP::HIS3/ REC8-EGFP::HIS3, cin8Δ:: loxp-URA3-loxp/ cin8Δ:: loxp-URA3-loxp, kip3Δ:: loxp-KanMx-loxp/ kip3Δ:: loxp-KanMx-loxp* |
| SGY5501 | *MAT*a/α, *REC8-EGFP::HIS3/ REC8-EGFP::HIS3, cin8Δ:: loxp-URA3-loxp/ cin8Δ:: loxp-URA3-loxp,* |
| SGY5523 | *MAT*a/α, *REC8-EGFP::HIS3/ REC8-EGFP::HIS3, SPC42-CFP::TRP1/ SPC42-CFP::TRP1, cin8Δ:: loxp-URA3-loxp/ cin8Δ:: loxp-URA3-loxp, kip3Δ:: loxp-KanMx-loxp/ kip3Δ:: loxp-KanMx-loxp* |
| SGY5532 | *MAT*a/α, *REC8-6HA::HIS3/ REC8-6HA::HIS3, PDS1-13MYC::TRP1/ PDS1-13MYC::TRP1, cin8Δ:: loxp-URA3-loxp/ cin8Δ:: loxp-URA3-loxp* |
| SGY5533 | *MAT*a/α, *REC8-6HA::HIS3/ REC8-6HA::HIS3, PDS1-13MYC::TRP1/ PDS1-13MYC::TRP1, cin8Δ:: loxp-URA3-loxp/ cin8Δ:: loxp-URA3-loxp, kip3Δ:: loxp-KanMx-loxp/ kip3Δ:: loxp-KanMx-loxp* |
| SGY5534 | *MAT*a/α, *REC8-6HA::HIS3/ REC8-6HA::HIS3, PDS1-13MYC::TRP1/ PDS1-13MYC::TRP1* |
| SGY5539 | *MAT*a/α, *KIP1-6HA::HIS3/ KIP1-6HA::HIS3* |
| SGY5540 | *MAT*a/α, *KIP1-6HA::HIS3/ KIP1-6HA::HIS3, cin8Δ:: loxp-URA3-loxp/ cin8Δ:: loxp-URA3-loxp, kip3Δ:: loxp-KanMx-loxp/ kip3Δ:: loxp-KanMx-loxp* |
| SGY5546 | *MAT*a/α, *leu2::tetR-GFP::LEU2, CEN5::TetO-HIS3/leu2::tetR-GFP::LEU2 CEN5::TetO-HIS3, SPC42-mCherry::KanMx/ SPC42-mCherry::KanMx, cin8Δ:: Loxp/ cin8Δ:: Loxp, kip3Δ:: Loxp/ kip3Δ:: Loxp, (*pJKY2*::URA3)* |

Santanu K Ghosh

| | |
|---|---|
| SGY5547 | MATa/α, *leu2::tetR-GFP::LEU2, CEN5::TetO-HIS3/leu2::tetR-GFP::LEU2 CEN5::TetO-HIS3, SPC42-mCherry::KanMx/ SPC42-mCherry::KanMx, cin8Δ:: Loxp/ cin8Δ:: Loxp, kip3Δ:: Loxp/ kip3Δ:: Loxp,* (pJKY12*::URA3)* |
| SGY5557 | MATa/α, *REC8-EGFP:: HIS3/ REC8-EGFP:: HIS3, SPC42-CFP::TRP1/ SPC42-CFP::TRP1* |
| SGY5561 | MATa/α, *KIP1-EGFP::TRP1/ KIP1-EGFP::TRP1 , SPC42-mCherry::KanMx/ SPC42-mCherry::KanMx, NDC80-CFP::HIS3/ NDC80-CFP::HIS3, kip3Δ:: Loxp, cin8Δ:: Loxp* |
| SGY5567 | MATa/α, *PDS1-EGFP::TRP1/ PDS1-EGFP::TRP1, cin8Δ:: loxp-URA3-loxp/ cin8Δ:: loxp-URA3-loxp, kip3Δ:: loxp-KanMx-loxp/ kip3Δ:: loxp-KanMx-loxp* |
| SGY5570 | MATa/α, *PDS1-EGFP::TRP1/ PDS1-EGFP::TRP1* |
| SGY5629 | MATa, *SPC42-mCherry::KanMx, MCD1-EGFP::HIS3* |
| SGY5630 | MAT α, *SPC42-mCherry::KanMx, kip3Δ:: Loxp, cin8Δ:: Loxp, MCD1-EGFP::HIS3* |
| SGY5634 | MATa/α, *SGO1-6HA::HPHMX/ SGO1-6HA::HPHMX, cin8Δ:: loxp-URA3-loxp/ cin8Δ:: loxp-URA3-loxp, kip3Δ:: loxp-KanMx-loxp/ kip3Δ:: loxp-KanMx-loxp* |
| SGY5635 | MATa/α, *SGO1-6HA::HPHMX/ SGO1-6HA::HPHMX* |
| SGY5610 | MATa/α, *spo11Δ:: loxp-LEU2-loxp/ spo11Δ:: loxp-LEU2-loxp, REC8-EGFP:: HIS3/ REC8-EGFP:: HIS3, SPC42-CFP:: TRP1/ SPC42-CFP:: TRP1* |
| SGY5628 | MATa/α, *mad2::*$P_{CLB2}$*-MAD2::URA3/ mad2::*$P_{CLB2}$*-MAD2::URA3 , REC8-EGFP::TRP1/ REC8-EGFP:: TRP1, SPC42-CFP:: HIS3/ SPC42-CFP:: HIS3* |
| SGY5667 | MATa/α, *rec8Δ:: loxp- HPHMX -loxp/ rec8Δ:: loxp-HPHMX -loxp* |
| SGY5670 | MATa/α, *leu2::tetR-GFP::LEU2, CEN5::TetO-HIS3/leu2::tetR-GFP::LEU2, CEN5::TetO-HIS3, cin8Δ:: loxp-URA3-loxp/ cin8Δ:: loxp-URA3-loxp, kip3Δ*:: *Loxp/ kip3Δ*:: *Loxp, rec8Δ:: loxp- HPHMX -loxp/ rec8Δ:: loxp-HPHMX -loxp* |
| SGY9002 | MATa/α, *leu2::tetR-GFP::LEU2, CEN5::TetO-HIS3/leu2::tetRGFP::LEU2, CEN5::TetO-HIS3, SPC42-mCherry::KanMx/ SPC42-mCherry::KanMx* |

Santanu K Ghosh

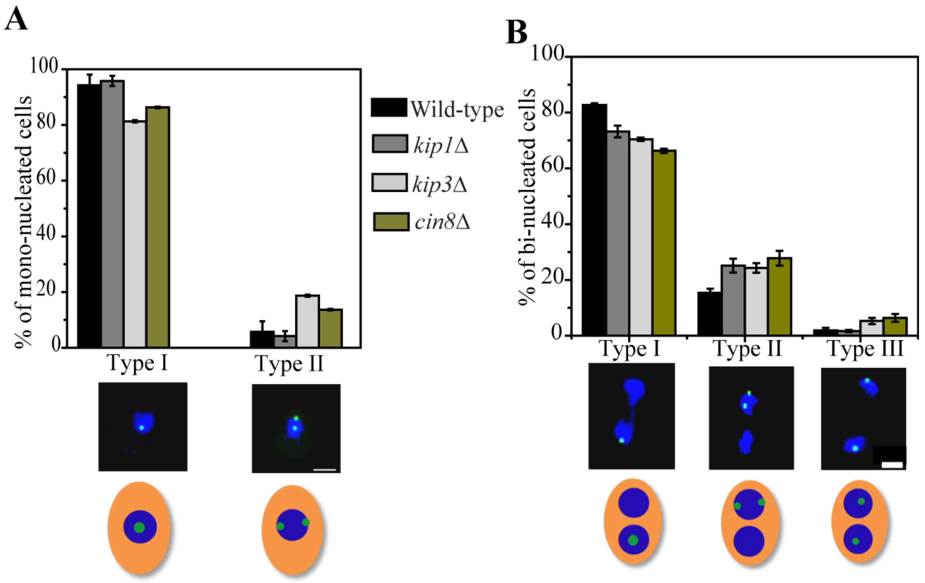


Figure S1: **Motor proteins do not have a role in the sister chromatid mono-orientation or cohesion.** Heterozygously marked CenV-GFP dots were analyzed for the meiotic chromosome segregation at 33°C. (A) Metaphase I arrested wild-type (SGY5143; n = 90), *cin8*Δ (SGY5078; n = 124), *kip1*Δ (SGY5018; n = 91) and *kip3*Δ (SGY5034; n = 71) cells were analyzed for the percentage of mono-nucleated with one or separated sister centromeres (n = 70 – 120). (B) Wild-type (SGY5006; n = 95), *cin8*Δ (SGY5090; n = 122), *kip1*Δ (SGY5077; n = 97) and *kip3*Δ (SGY309; n = 96) cells were analyzed for the percentage of bi-nucleates with one or two GFP dots in one nucleus (Type I and II, respectively) or one dot each in two nuclei (Type III). Error bars represent the standard deviation from the mean values obtained from three independent experiments and 'n' represents the number of cells scored. Bar, 2 µm.

Santanu K Ghosh

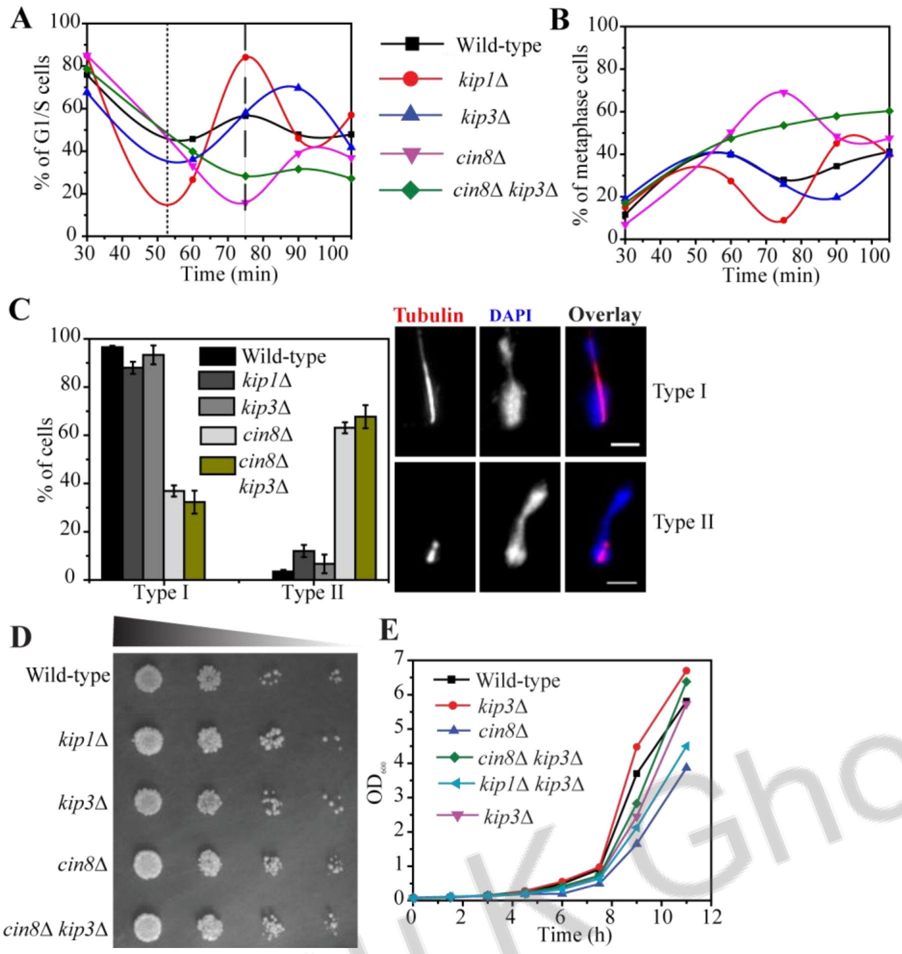


Santanu K Ghosh

Figure S2: ***cin8Δ kip3Δ* delays the metaphase to anaphase transition during mitosis but the mitotic growth is not affected.** (A and B) For synchronization, the wild-type (SGY155), *cin8Δ* (SGY5357), *kip1Δ* (SGY317), *kip3Δ* (SGY304), and *cin8Δ kip3Δ* (SGY5124) cells were arrested at G1 stage with α-factor and were released subsequently into YPD medium. Cells at different mitotic stages were determined by tubulin immunostaining in > 100 cells for each indicated time point. Dotted and dashed lines represent the time-points at the start of the second peak of G1/S stage cells in wild-type, *kip1Δ*, *kip3Δ* and in *cin8Δ*, *cin8Δ kip3Δ*, respectively. (C) Representative images of two types (Type I and Type II) of spindle morphologies within extended DAPI masses are shown on the right. The distribution of these types among the indicated strains is shown on the left. Error bars represent the standard deviation from the mean values obtained from three independent experiments. Bar, 2 µm. (D) 10-fold serial dilutions of 1 O.D$_{600}$ culture of the indicated strains were spotted on nutrient-rich YPD media and kept for incubation at 30°C for approximately 36 h before being photographed. (E) Growth curve of the same strains at 30° C. As shown the growth rate of *cin8Δ kip3Δ* is not significantly different from the wild-type.

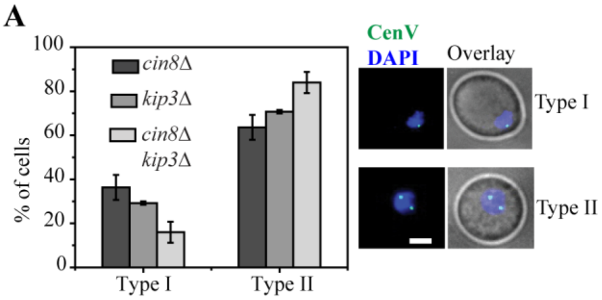


B

Percentage of cells

| | G1/S | | Metaphase | | Anaphase |
|---|---|---|---|---|---|
| | Type I | Type II | Type I | Type II | |
| Wild-type | 100 | 0 | 80 | 20 | 100 |
| *kip3Δ* | 95.2 | 4.8 | 79 | 21 | 100 |
| *cin8Δ* | 94.6 | 5.4 | 81.8 | 18.2 | 100 |
| *cin8Δ kip3Δ* | 95.6 | 4.4 | 65.4 | 34.6 | 100 |

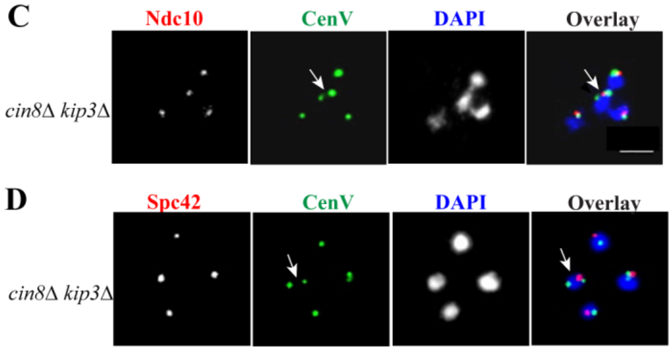


Figure S3: **Chromosome V aneuploidy in *cin8Δ kip3Δ* cells does not occur in mitosis, and in meiosis it occurs not due to chromosome misduplication or multipolarity.** (A-B) Supernumerary CenV-GFP is not generated during mitosis. (A) The *cin8Δ* (SGY315), *kip3Δ* (SGY314), and *cin8Δ kip3Δ* (SGY5089) cells with homozygous CenV-GFP were synchronized before meiosis induction in YPA (Yeast extract Potassium Acetate) and at the time of their release into the sporulation medium, the cells were analyzed for the number of GFP dots in at least 100 cells. (B) Segregation of CenV-GFP dots was analyzed at different stages of mitosis in the wild-type (SGY155), *cin8Δ* (SGY302), *kip3Δ* (SGY304), and *cin8Δ kip3Δ* (SGY5124) haploid cells ( n = 50-120). None of the cells with more than 2 GFP dots was observed. (C-D) Localization of supernumerary CenV-GFP foci along with the kinetochore (Ndc10) and spindle pole (Spc42). Unlike CenV-GFP, no cell showed more than four Ndc10-mcherry (SGY5333) (C) or Spc42-mcherry (SGY5338) (D) foci in the *cin8Δ kip3Δ* cells. Arrows indicate a single Ndc10 or Spc42 focus between 2 CenV-GFP foci. Error bars represent the standard deviation from the mean values obtained from three independent experiments. Bar, 2 µm.

Santanu K Ghosh

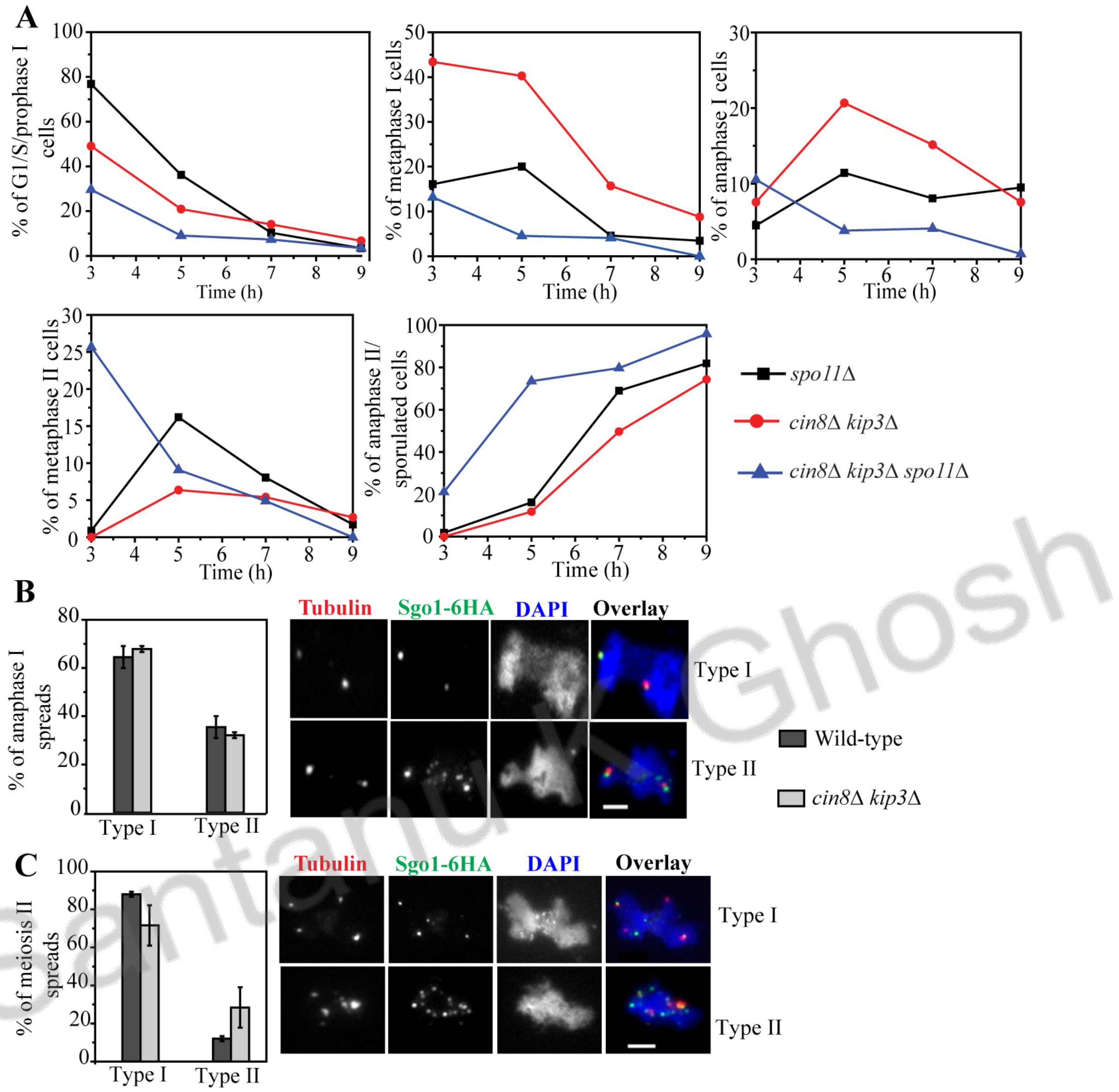


Santanu K Ghosh

Figure S4: **The anaphase I delay observed in *cin8*Δ *kip3*Δ due to persistent cohesin on the chromatin is relieved by inhibition of recombination but the persistence is not mediated by Sgo1.** (A) The *spo11*Δ (SGY5048), *cin8*Δ *kip3*Δ (SGY5089) and *cin8*Δ *kip3*Δ *spo11*Δ (SGY5444) cells were analyzed for meiotic progression where different stages of meiosis were judged by tubulin immunostaining and DAPI staining at the indicated time points. At least 90 cells were scored for each time point. (B-C) The chromatin spreads from the wild-type (SGY5635; n = 110), and *cin8*Δ *kip3*Δ (SGY5634; n = 128) cells were analyzed for Sgo1-6HA localization during (B) bi-nucleated and (C) meiosis II stages of meiosis using anti-HA and anti-tubulin antibodies. ‘n’ represents the total number of chromosome spreads examined. Error bars represent the standard deviation from the mean values obtained from three independent experiments. Bar, 2μm.
